# Disruption of a GATA2, TAL1, ERG regulatory circuit promotes erythroid transition in healthy and leukemic stem cells

**DOI:** 10.1101/2020.10.26.353797

**Authors:** Julie A. I Thoms, Kathy Knezevic, Gregory Harvey, Yizhou Huang, Janith A. Seneviratne, Daniel R. Carter, Shruthi Subramanian, Joanna Skhinas, Diego Chacon, Anushi Shah, Ineke de Jong, Dominik Beck, Berthold Göttgens, Jonas Larsson, Jason W. H. Wong, Fabio Zanini, John E. Pimanda

## Abstract

Changes in gene regulation and expression govern orderly transitions from hematopoietic stem cells to terminally differentiated blood cell types. These transitions are disrupted during leukemic transformation but knowledge of the gene regulatory changes underpinning this process is elusive. We hypothesised that identifying core gene regulatory networks in healthy hematopoietic and leukemic cells could provide insights into network alterations that perturb cell state transitions. A heptad of transcription factors (LYL1, TAL1, LMO2, FLI1, ERG, GATA2, RUNX1) bind key hematopoietic genes in human CD34+ haematopoietic stem and progenitor cells (HSPCs) and have prognostic significance in acute myeloid leukemia (AML). These factors also form a densely interconnected circuit by binding combinatorially at their own, and each other’s, regulatory elements. However, their mutual regulation during normal haematopoiesis and in AML cells, and how perturbation of their expression levels influences cell fate decisions remains unclear. Here, we integrated bulk and single cell data and found that the fully connected heptad circuit identified in healthy HSPCs persists with only minor alterations in AML, and that chromatin accessibility at key heptad regulatory elements was predictive of cell identity in both healthy progenitors and in leukemic cells. The heptad factors GATA2, TAL1 and ERG formed an integrated sub-circuit that regulates stem cell to erythroid transition in both healthy and leukemic cells. Components of this triad could be manipulated to facilitate erythroid transition providing a proof of concept that such regulatory circuits could be harnessed to promote specific cell type transitions and overcome dysregulated haematopoiesis.

## INTRODUCTION

Haematopoietic stem cells (HSCs) reside in the bone marrow niche where they are mostly quiescent but retain the capacity to self-renew and replace terminal blood cell types throughout life^1^. Haematopoiesis is a hierarchical process with HSCs at the apex giving rise to a range of progenitor cells with increasing lineage restriction^1^. Although single cell transcriptomic data suggest a rather continuous differentiation process^2–7^, relatively pure progenitor populations corresponding to intermediate differentiation stages can be prospectively isolated based on surface marker expression^3^. Cell type transitions are controlled by cell intrinsic and extrinsic factors, and loss of control can lead to inappropriate proliferation and leukemic transformation^8–13^.

Acute myeloid leukemia (AML) is characterised by an abundance of relatively undifferentiated cells (blasts) of the myeloid lineage^14^. AMLs likely originate in the earliest HSC compartments or acquire stem-cell-like transcriptional programs during leukemic transformation^15–19^. Although blast cells can comprise the bulk of the AML population, self-renewal is restricted to a smaller population of leukemic stem cells (LSCs) which can recapitulate the disease after ablation of the blast population^20–22^. LSCs drive relapse^23^, potentially because they possess stem cell transcriptional programs^24,25^. Thus, AML induces a parallel hierarchy of malignant cell types with LSCs at the top^26^. Therapies that induce LSC differentiation by targeting mutant proteins that block differentiation are effective but limited to a minority of AMLs^27–31^.

AML is a heterogenous disease with many known driver mutations^14,32–34^, however it is clear that many of these drivers converge on corruption of the transcriptional networks that control normal haematopoiesis^13,35–37^. Transcriptional networks coordinate gene regulation and play a key role in establishing and maintaining cell identity throughout the life of an organism^12,38^. Such networks are cell type specific, and therefore need to be rewired during embryonic development and differentiation, while their disruption can lead to oncogenic transformation^8–13^. Indeed, transcriptional networks are altered across AMLs with a wide spectrum of mutational origins, such that AML cells assume a new epigenetic identity which is distinct from any normal blood cell type^35^. Furthermore, epigenetic rewiring is increasingly being recognised as a non-genetic cause of treatment resistance^39–41^. However, the specific molecular mechanisms underlying disruption of transcriptional networks in AML, and whether these can be therapeutically targeted, remain unknown.

We and others have previously described seven transcriptional regulators (heptad; LYL1, TAL1, LMO2, FLI1, ERG, GATA2, RUNX1) which bind to key haematopoietic genes in normal human CD34+ haematopoietic stem and progenitor cells (HSPCs) and in AML^42–44^. Heptad factors also bind combinatorially at their own, and each other’s, regulatory elements, forming a densely interconnected circuit^42,44^. The heptad circuit appears to be established at the haemogenic endothelium stage of blood development^45^, and over-expression of all seven factors in a mouse *in vitro* differentiation system led to increased production of pre-HSPCs with capacity for multilineage differentiation ^46^. All seven factors are key haematopoietic regulators, and mutation or dysregulation is commonly associated with haematological or other malignancies^32,47–50^. Furthermore, the heptad circuit is maintained or reactivated in AML^43,51–53^, and heptad expression is predictive of patient outcome^43^. However, heptad circuitry and function have primarily been established using bulk ChIPseq experiments in heterogenous cell populations (i.e. CD34^+^ HSPCs) which may obscure underlying sub-circuits or relationships that only exist in specific cell types/cellular contexts. Thus, key questions remain about the precise role of the heptad throughout normal and leukemic haematopoiesis, including whether all seven factors act together in single cells, and whether heptad perturbation can influence cell fate decisions.

Here we integrate bulk and single cell data in normal human HSPCs and leukemic cells and find that chromatin conformation at key heptad regulatory elements is predictive of cell identity in normal and leukemic progenitors. The interconnected heptad circuit identified in normal HSPCs persists in AML, but single cell transcriptomics suggest that specific heptad sub-circuits exist in individual cells.

## METHODS

Supplementary Methods have extended details of standard techniques.

### Cell culture and NGS data generation

Cells were cultured, and luciferase reporter assays performed, using standard techniques. Chromatin immunoprecipitation (ChIP) was performed as described^43^ (antibodies in Table S1). Library construction/sequencing was performed by BGI Genomics (China) or Novogene (Hong Kong). Single cell RNA sequencing (scRNAseq) data was generated using the 10X Genomics pipeline.

### NGS data processing

ATAC, DHS, and ChIPseq data was aligned and displayed in BigWig format, and peaks called using standard analysis pipelines. RNAseq data was processed using standard techniques. Total read counts covering enhancer regions (Table S2) were extracted using pyBigWig (https://github.com/deeptools/pyBigWig) and plotted.

For normal cell ATACseq, counts from replicate experiments were added together before plotting. Each profile was encoded as a unit vector by dividing by the total counts across all studied heptad peaks. Cityblock distances on the multidimensional unit sphere between each sample and each average profile were used to compute the heatmap and predict cell types.

### sc-RNAseq Analysis

The secondary analysis for Figures 1 and 4 is included at https://github.com/iosonofabio/heptad_paper. Single cell RNA-Seq data on healthy hematopoietic cells was downloaded from the Palantir github repository as described on https://github.com/dpeerlab/Palantir/blob/master/README.md, Rep1. Embedding coordinates, colours, cluster metadata, and smoothed counts data were extracted from the h5ad file and plotted using singlet (https://github.com/iosonofabio/singlet).

**Figure 1.**
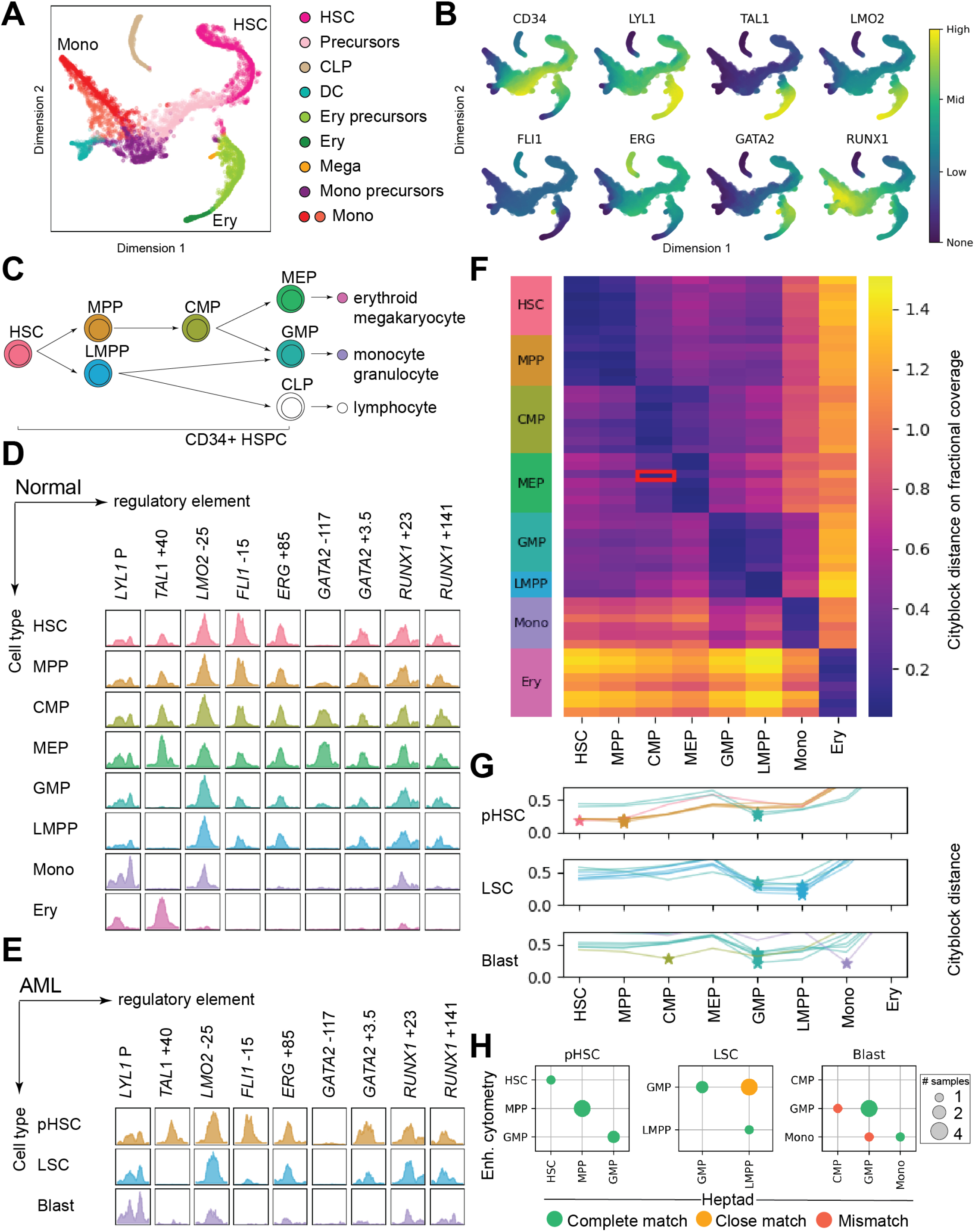
Heptad regulatory regions have dynamic accessibility profiles across normal and leukemic blood development, and accessibility patterns are sufficient to classify normal and leukemic cells. (A) tSNE plot of single cell RNAseq in normal bone marrow, with cells labelled by inferred identity as determined by Setty et al 2019. HSC = haematopoietic stem cell, CLP = common lymphoid progenitor, DC = dendritic cell, Ery = erythoid lineage cells, Mega = megakaryocytes, Mono = monocyte lineage cells. (B) Relative expression of CD34 and heptad genes projected on to the tSNE plot in A. (C) Schematic of the branching hierarchy model of normal blood development showing relationships between the cell populations shown in D. (D) ATACseq peaks at heptad regulatory regions over developmental time. Plots show merged data from available replicates (Corces et al 2016). (E) ATACseq peaks at heptad regulatory regions in one representative AML patient, showing pre-leukemic HSCS (pHSC), leukemic stem cells (LCS), and leukemic blasts (Blast). (F) Classification of normal cell types using only ATACseq signal at heptad regulatory regions. Heatmap shows calculated distance between each sample and the training set. The red box indicates a single MEP replicate that was misclassified as a CMP. (G) Classification of AML nearest normal cell type using only ATACseq signal at heptad regulatory regions. Plots show distance from each normal cell type for pre-leukemic HSCS, LSCs, and leukemic blasts from seven AML patients. (H) Performance of the heptad regulatory region classifier compared to previous classification of these samples using genome wide enhancer cytometry (Corces et al 2016).

Count and metadata tables from CellRanger (10X Genomics) were converted to loom format (http://loompy.org/) and normalised to “counts per ten thousand (uniquely mapped) reads” The symmetric correlation matrix was ordered by hierarchical (average linkage) clustering on L2 distance with optimal leaf ordering. Conditional distributions of gene expression were computed via quantiles followed by kernel density estimate in logarithmic space.

Palantir data were subsampled to 40 cells per cell type. northstar’s subsample method^54^ was used to infer cell states within ME-1 guided by Palantir data^6^. In the graph construction step, 10 external (non-mutual) neighbours were allowed in order to compensate for the fact that ME-1 cells are quite distant from actual hematopoietic cells. ME-1 cells were then coloured based on northstar’s community assignments. RNA velocity^55^ was computed using scVelo^56^ and projected onto northstar’s embedding. Gene expression was plotted in the same embedding after iterative nearest-neighbour smoothing. For the prediction of ME-1 cell state based on heptad and subsets thereof, we trained a random forest classifier using scikit-learn and evaluated its performance via train/test splits.

### Data sharing statement

Published datasets used are listed in Table S3. Data generated here is available using GEO accession GSE158797. The code is available from https://github.com/iosonofabio/heptad_paper.

## RESULTS

### Heptad expression during haematopoiesis

To understand patterns of heptad expression during haematopoiesis we interrogated existing single cell RNAseq data (Palantir) from bone marrow cells^6^ (Figure 1A). Diverging patterns of heptad transcription factor (TF) expression were observed across developmental time (Figure 1B). All seven TFs are expressed in HSCs, with increasing divergence during differentiation. For example, *GATA2*, *TAL1*, *LYL1*, and *LMO2* are upregulated along the erythroid lineage, while *RUNX1* is upregulated along the granulocytic/monocytic lineage.

### Heptad regulatory region accessibility during normal haematopoiesis

Heptad TFs form a densely interconnected circuit in bulk CD34+ HSPCs, with each corresponding gene having regulatory regions bound by most of the heptad^42^. Since heptad expression patterns are heterogeneous in single cells, we asked whether there is evidence for changes in heptad regulation at any of the combinatorially bound regions over developmental time. Although haematopoiesis is a continuum (Figure 1A), functionally defined subpopulations representing various waypoints can be isolated based on cell surface marker expression (Figure 1C). We queried chromatin accessibility data from sorted bone marrow subpopulations^4^, focussing on known heptad gene regulatory regions (*LYL1* promoter (P), *TAL1* +40, *LMO2* −25, *FLI1* −16, *ERG* +85, *GATA2* +3.5, *RUNX1* +23^42^). We included two putative regulatory regions; *RUNX1* +141, an intragenic *RUNX1* region that was heptad-bound in CD34+ HSPCs^42^, and *GATA2* −117, a distal regulatory element for *GATA2* that is dysregulated by translocation in the inv(3) AML subtype^57,58^. Strikingly, accessibility patterns differed throughout development with elements losing accessibility upon exiting the CD34^+^ progenitor stage, suggesting that heptad connectivity is lost once cells commit to terminal differentiation (Figure 1D). Changes in chromatin accessibility reflected gene expression (compare Figures 1B, 1D). For example, *TAL1* expression increased as cells progressed towards erythroid lineages and *TAL1* +40 accessibility was restricted to the HSC to erythroid axis. Likewise, *GATA2* expression (highest in erythroid precursors and reduced in mature erythroid cells) was reflected by accessibility of the *GATA2* −117 element.

### Heptad regulatory region accessibility in AML

The heptad circuit can be active in AML^43,51–53^ and heptad expression can predict patient survival^43^. Data from two cohorts of AML cells showed that heptad regulatory regions were accessible in AMLs with diverse molecular lesions^35^ (Figure S1A) and in pre-leukemic HSCs, LSCs, and leukemic blasts isolated from the same patient^4^ (Figures 1E, S1B). Notably, the *TAL1* +40 enhancer was rarely accessible in AML, and the *GATA2* −117 enhancer varied between patient samples.

### Heptad regulatory region accessibility can classify normal and leukemic cells

Genome-wide chromatin accessibility profiles reflect cell identity^4^. Since heptad expression and regulatory region accessibility are restricted to progenitor populations in normal blood, and heterogenous throughout development, we asked whether the pattern of chromatin accessibility at heptad regulatory regions is sufficient to predict cell type. Using a classifier based on nine regulatory regions, we could correctly identify normal cells across the haematopoietic spectrum (Figure 1F). Furthermore, this classifier could assign a “closest normal” type to AML samples sorted into pre-leukemic HSC (pHSC), LSC, and blast populations (Figure 1G). Consistent with known AML biology, pHSCs were predominantly classified as HSCs or MPPs, LSCs as LMPPs or GMPs, and blasts as more variable cell types. We compared our cell type assignments to published classifications of these samples based on whole genome accessibility patterns^4^ and found a high concordance in pHSC and LSC populations (Figures 1H, S1C).

Consistent with the loss of heptad connectivity in more differentiated cells, the heptad-based classifier had reduced concordance with genome-wide classification in blast populations. Overall, our analysis indicates that heptad expression and accessibility are associated with cell identity in healthy haematopoietic progenitors and leukemic cells.

### The heptad network persists in AML, with altered connectivity

We extended our analysis and asked which heptad TFs were bound at each regulatory region in normal and AML contexts, looking first at heptad binding patterns at the nine regulatory regions in CD34^+^ HSPCs^42^ (Figure 2A, *left*, Figure S2). Combinatorial binding was observed with LYL1, FLI1, GATA2, and RUNX1 bound at all regions, and *FLI1*, *ERG*, *GATA2*, and *RUNX1* each having at least one regulatory element bound by all seven TFs. Binding patterns were then used to infer the connectivity map of heptad autoregulation in CD34+ HSPCs (Figure 2A, *right*).

**Figure 2.**
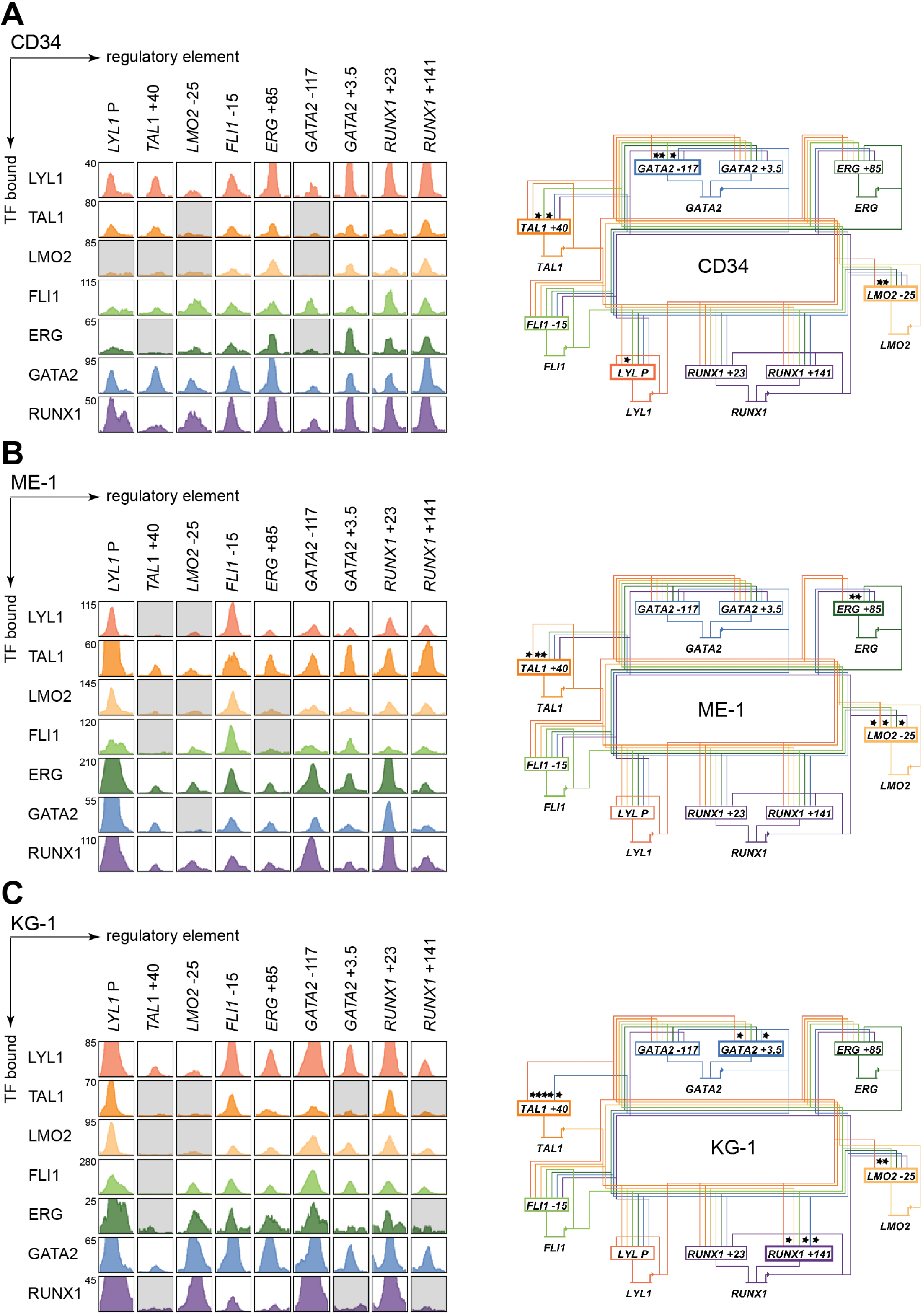
A densely interconnected heptad autoregulatory circuit persists in AML cells with altered connectivity compared to CD34+ HSPCs. (A) *Left:* ChIPseq binding pattern at heptad regulatory regions in CD34^+^ HSPCs. Grey boxes indicate regulatory regions not computationally called as binding peaks for the indicated TF. Plots are scaled to 5x the height of the smallest called peak for that TF to allow visualisation of a wide range of peak heights. *Right:* Corresponding inferred heptad autoregulatory circuit. Most regulatory elements have all seven heptad TFs bound, * and bold border indicate regions where binding of a particular TF is absent. (B) *Left:* ChIPseq binding pattern at heptad regulatory regions in ME-1 AML cells. Grey boxes indicate regulatory regions not computationally called as binding peaks for the indicated TF. Plots are scaled to 5x the height of the smallest called peak for that TF to allow visualisation of a wide range of peak heights. *Right:* Corresponding inferred heptad autoregulatory circuit. Most regulatory elements have all seven heptad TFs bound, * and bold border indicate regions where binding of a particular TF is absent. (C) *Left:* ChIPseq binding pattern at heptad regulatory regions in KG-1 AML cells. Grey boxes indicate regulatory regions not computationally called as binding peaks for the indicated TF. Plots are scaled to 5x the height of the smallest called peak for that TF to allow visualisation of a wide range of peak heights. *Right:* Corresponding inferred heptad autoregulatory circuit. Most regulatory elements have all seven heptad TFs bound, * and bold border indicate regions where binding of a particular TF is absent

We next compared heptad connectivity in two AML cell lines, ME-1, and KG-1. AML cell lines recapitulate properties of primary AML cells^59^ and are amenable to experimental manipulation. Consistent with accessibility in primary AMLs, heptad ChIPseq in ME-1 (Figures 2B, S3) and KG-1 (Figure 2C, S4) revealed that the densely interconnected circuit observed in CD34^+^ HSPCs persists in AML cells. Notably, both ME-1 and KG-1 have binding peaks at the *LYL1* promoter, while at *TAL1* +40, ME-1 and KG-1 had fewer called peaks (4/7 and 2/7 respectively) than CD34+ HSPCs (5/7), and these were generally small. Overall, heptad connectivity is maintained in both AML cell lines, albeit with somewhat different connection patterns to those observed in HSPCs.

### Heptad regulatory elements require ETS and GATA motifs

Having shown that heptad binding at regulatory regions persists in AML, we wanted to understand the regulatory regions and the role of specific TF binding motifs within these elements. *Cis*-regulatory elements integrate signals from multiple TFs which bind to specific DNA sequences, with direct binding occurring at consensus binding motifs. The heptad TFs belong to four broad classes of TFs with different consensus binding motifs – E-box (CANNTG, bound directly by LYL1 and TAL1 and indirectly by LMO2), ETS (GGAW, bound by FLI1 and ERG), GATA (GATA, bound by GATA2) and RUNX (TGYGGT, bound by RUNX1). To identify consensus motifs likely to correspond to TF binding sites, we performed multiple sequence alignments using human, mouse, dog, and opossum genomes (Figure 3A). All regulatory elements contained conserved ETS and GATA motifs, while 7/9 contained a conserved E-Box motif and 6/9 a conserved RUNX motif. We mutated all conserved instances of each binding motif class (Table S4) and tested in luciferase reporter constructs in both KG-1 and ME-1 cells.

**Figure 3.**
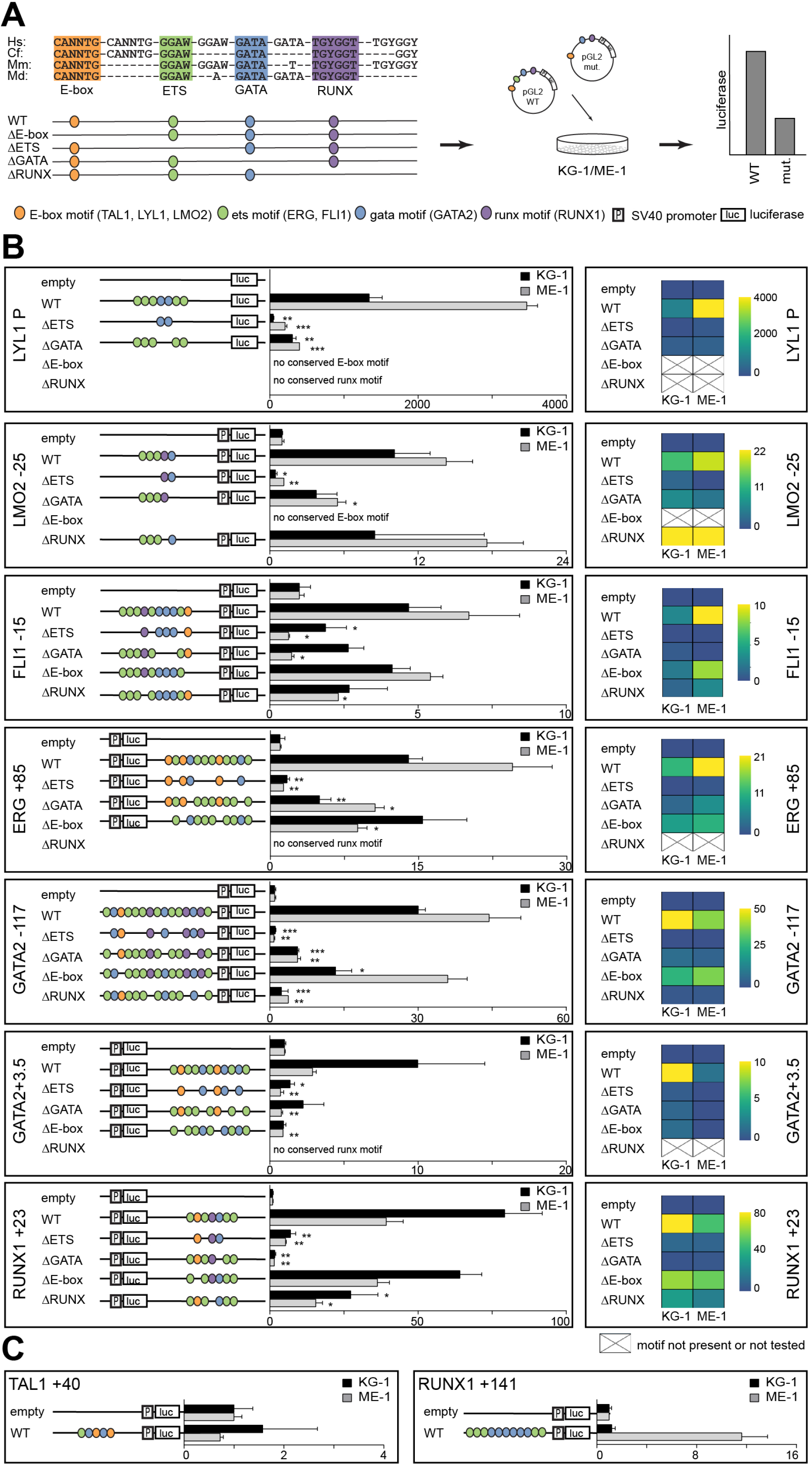
Specific TF consensus binding motifs, particularly ETS and GATA motifs, are critical for function of heptad regulatory elements. (A) Schematic showing process for selecting TF binding motifs for mutation, and luciferase reporter workflow. (B) *Left panel:* Schematics showing conserved TF binding motifs in heptad regulatory elements that were highly bound by heptad TFs in AML cell lines, and activity of wild type (WT) and mutated luciferase constructs in KG-1 and ME-1 cells. Activity is scaled relative to the empty vector, and graphs show representative data from a single transfection experiment (* P < 0.05, ** P < 0.01, *** P < 0.001, t-test). *Right panel:* Heatmaps showing aggregate data from all luciferase experiments. Data from biological replicates were normalised to WT activity for each experiment, then aggregate data scaled relative to empty vector. Heatmaps are scaled from 0 to maximum luciferase activity for each regulatory element. (C) Schematics showing conserved TF binding motifs in heptad regulatory elements that were highly bound by heptad TFs in AML cell lines, and activity of WT luciferase constructs in KG-1 and ME-1 cells. Activity is scaled relative to the empty vector, and graphs show representative data from a single transfection experiment (* P < 0.05, ** P < 0.01, *** P < 0.001, t-test).

Deletion of ETS consensus motifs was universally deleterious, leading to significant loss of activity for all elements tested (Figure 3B). Deletion of GATA consensus motifs had a significant negative impact for all regions in at least one cell line. Deletion of E-box or RUNX motifs reduced luciferase reporter activity, however the effect was generally small compared to deletion of ETS or GATA motifs, and in one case (*LMO2* −25) deletion of the RUNX motif lead to slightly increased activity. Overall, regulatory region activity was impaired by loss of any class of TF binding motif, with loss of ETS or GATA motifs dominating. Two WT reporter constructs, *TAL1* +40 and *RUNX1* +141, showed minimal activity in one or both cell lines (Figure 3C), and were excluded from mutation analysis.

### Single cell transcriptomics reveal key regulators of the HSC – erythroid transition

Altered enhancer activity reads out as gene expression changes. Encouraged by our results indicating that removing specific consensus motifs altered activity of heptad regulatory regions, we proceeded to single cell RNAseq analysis of heptad expression. For this analysis we selected ME-1 cells which are amenable to downstream perturbation. We quantified heptad heterogeneity and observed that for both high (e.g. *LYL1*) and low (e.g. *ERG*) expressed genes heterogeneity across the ME-1 population spanned an order of magnitude (Figure 4A). Furthermore, the highest gene expression (*LYL1*) corresponded to the highest heptad binding at an associated regulatory region, while lower gene expression (*TAL1* and *GATA2*) corresponded to lower heptad binding at their associated regulatory regions (Figure 2B).

**Figure 4.**
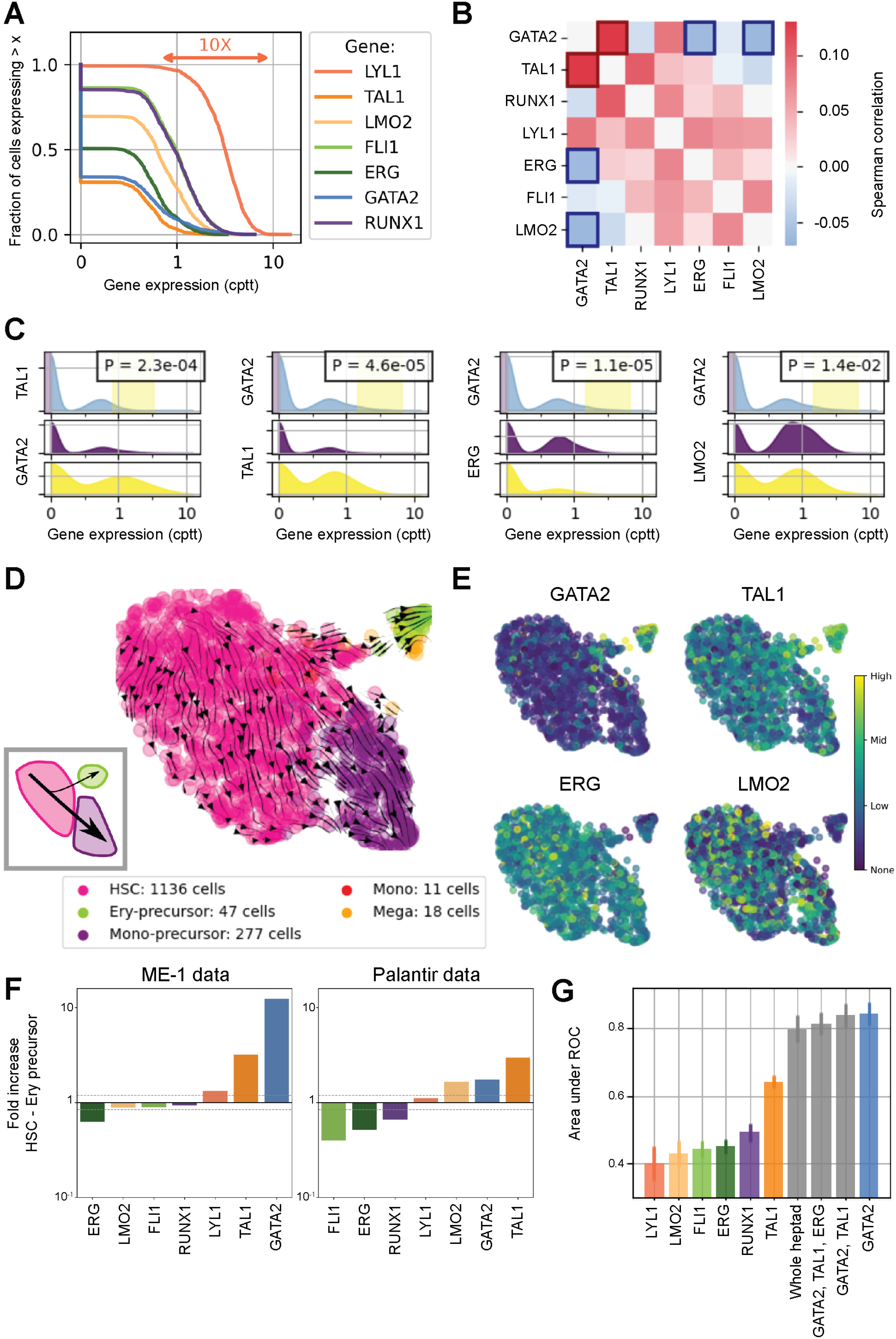
Single cell transcriptomics in ME-1 cells reveals branching heterogeneity consistent with GATA2 regulation. (A) Cumulative expression distributions for heptad genes in single ME-1 cells. cppt: counts per ten thousand reads. (B) Pairwise Spearman correlations between heptad genes in single cells. (C) Censored distributions of gene expression for the gene pairs highlighted in B. The two lower panels show the expression of the second gene in the lowest 10% and highest 5% of expressing cells for the first gene. P values refer to a Kolmogorov-Smirnov 2-sample test between the purple and yellow distributions. (D) UMAP embedding of ME-1 cells and cell state assignment based on northstar (Zanini et al 2020) and the Palantir data as atlas (see Figure 1). Stream-lines show RNA velocity as computed by scVelo (Bergen et al 2019), projected onto the same embedding. *Inset:* Schematic of the branching phenotype within ME-1 cells, indicating the cell flux into the Ery-precursor-like state is a rare event. (E) Expression of four heptad genes highlighted in B on the embedding. Colour legend: purple = no expression, green = low expression, yellow = high expression. (F) *Left:* Fold increase in heptad gene expression across the HSC to Ery-precursor-like state in ME-1 cells. *Right:* Fold increase in heptad gene expression across the HSC to Ery-precursor state in normal CD34^+^ HSPCs cells. (G) Performance of random forest classifiers between HSC-like and Ery-precursor-like states in ME-1 trained solely on Palantir data with a spectrum of selected features. The presence of GATA2 expression in the model is essential for its accuracy. Error bars indicate SD over 10 runs of the predictor with data resampling in each run.

We next asked whether there were any pairwise expression correlations between TFs and found that *GATA2* was positively correlated with *TAL1*, and negatively correlated with *ERG* and *LMO2* (Figure 4B). Because correlation measures are insensitive to extreme phenotypes, we performed complementary analysis to evaluate whether this effect is also seen at the extreme of the distribution and plotted conditional gene expression distribution in the bottom and top quantiles of expressors of *GATA2* (Figure 4C). Given the observed heterogeneity in heptad expression in ME-1 cells, and the strong association between heptad regulation and cell type we asked whether we could identify subpopulations within the ME-1 sc-RNAseq data. A canonical unsupervised clustering approach based on overdispersed features did not result in distinct biological patterns beyond cell cycle, as expected from a cell line. We reasoned that a more sophisticated feature selection together with soft guidance from healthy marrow data could reveal additional hidden heterogeneity. We therefore switched from unsupervised clustering to northstar, a semi-supervised clustering algorithm that leverages information from training data to channel the axes of heterogeneity during feature selection, graph construction, and cell community detection^54^. Using healthy marrow transcriptomes^6^ (Figure 1A) as training data, this analysis revealed two major subpopulations, HSC-like (pink) and Mono-precursor-like (purple, 1136 and 277 out of 1489 cells respectively) plus a minor population that was more similar to Ery-precursor cells (lime, 47 out of 1489 cells) and two small groups of cells resembling Megakaryocytes (18 cells) and Monocytes (11 cells, Figure 4D). RNA velocity analysis^55^ (Figure 4D *arrows*) revealed a major trajectory along the HSC-Mono-precursor axis, and an alternate trajectory connecting the HSCs to the Ery-precursor population. This flow diagram (independent of northstar clustering) confirmed population structure reminiscent of healthy haematopoiesis (Figure 4D *inset*). We next projected expression levels of the four previously identified genes on embedded cell plots (Figure 4E). Consistent with our correlation data and known biological functions, *GATA2* and *TAL1* expression were enriched in the Ery-precursor population. Conversely, *ERG* and *LMO2* expression were enriched in the HSC-like and Mono-precursor-like populations. We then computed the fold expression change in heptad genes between HSC and Ery-precursor cells in both ME-1 and normal BM cells (Figures 4F, S5, Tables S5, S6). In ME-1 cells, *ERG* expression was reduced (0.6x) and *GATA2* and *TAL1* expression increased (11x and 3.5x respectively) in Ery-precursor cells (Figure 4F *left*). We observed a similar pattern in healthy cells, although *FLI1*, *RUNX1*, and *LMO2* also showed expression changes in this context (Figure 4F *right*).

Finally, we asked whether heptad expression was sufficient to classify ME-1 cells as HSC-like or Ery-Precursor-like (Figure 4G). Using a random forest classifier based on Palantir data, we found heptad expression was able to correctly classify cells with high accuracy (area under ROC = 0.80), and that *GATA*2 expression was the single best performing gene in terms of model accuracy (area under ROC = 0.84).

### Direct manipulation of GATA2 and ERG promotes erythroid trajectory

We then evaluated the effect of perturbing key heptad factors on i) expression of other heptad factors, and ii) the global transcriptome of perturbed cells. Specifically, we predicted that high levels of *GATA2* or *TAL1* and low levels of *ERG* would promote transition along the HSC-Ery-precursor axis (Figure 5A). We first knocked down key heptad genes in ME-1 cells and measured the response of other heptad genes. Knockdown of *GATA2* led to decrease of *TAL1* and most other heptad genes, with the exception of *ERG* which was unaffected by *GATA2* knockdown (Figure 5B, *left*). Similarly, knockdown of *TAL1* led to decreased *GATA2* and most other heptad genes except for *ERG* (Figure 5B, *centre*). Conversely, *ERG* knockdown led to decreased *LMO2* expression, but increased expression of *GATA2*, *FLI1*, and *TAL1* (Figure 5B, *right*). *RUNX1* expression did not change in a consistent way, possibly due to dysregulation via translocation of its essential binding partner *CBFβ* in ME-1 cells^60^.

**Figure 5.**
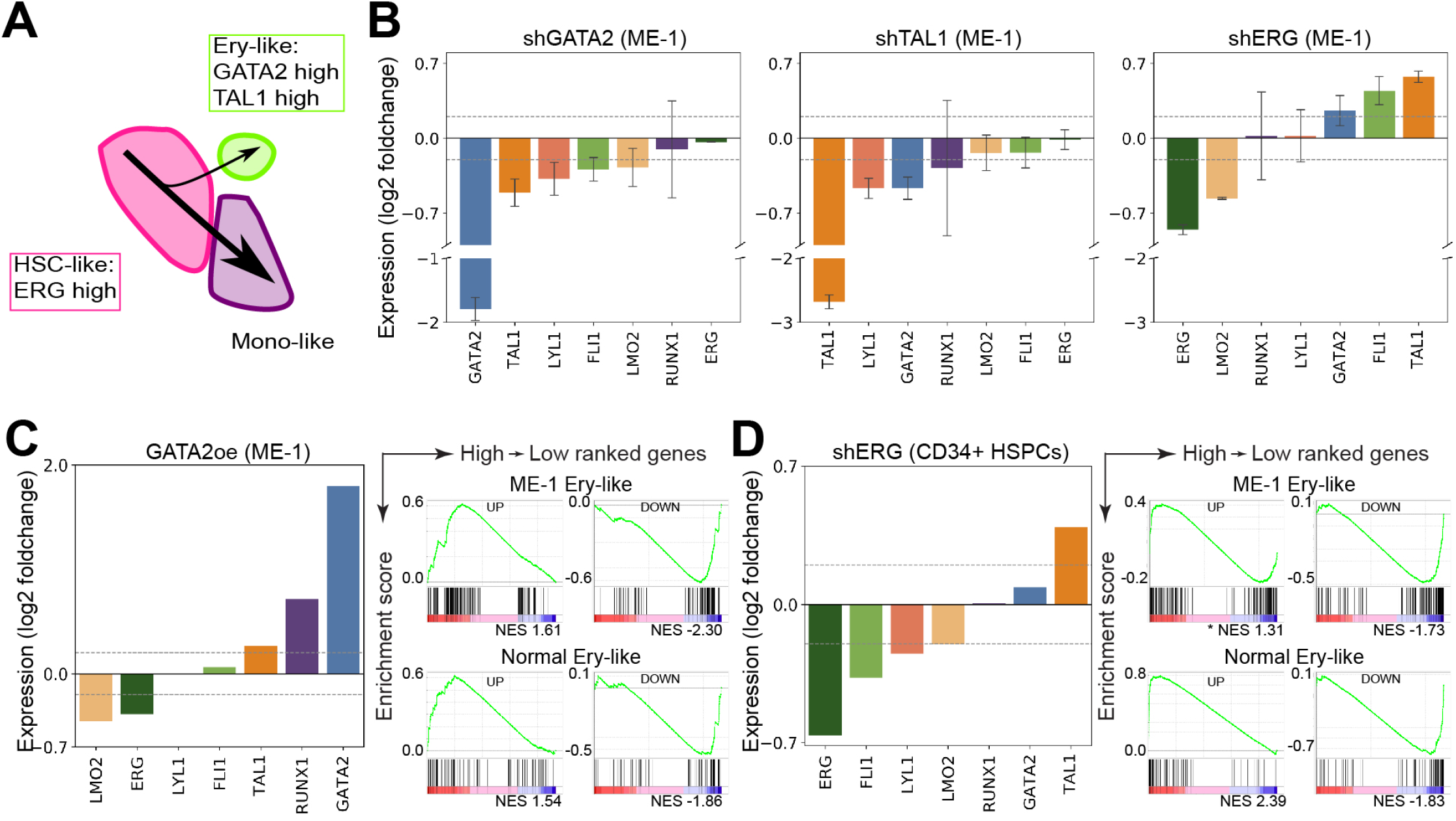
Manipulating GATA2 and ERG in bulk ME-1 cells and normal CD34+ HSPCs leads to altered heptad expression and can push cells towards the Ery-like state. (A) Schematic of the branching phenotype within ME-1 cells indicating relative expression of key heptad genes highlighted in Figure 4. (B) Effect of knocking down *GATA2*, *TAL1*, or *ERG* on heptad genes in ME-1 cells. (C) *Left:* Effect of over-expressing GATA2 on heptad genes in ME-1 cells. *Right:* GSEA plots showing enrichment of genes associated with the Ery-precursor/Ery-precursor-like state in response to over-expressing GATA2 in ME-1 cells. (D) *Left:* Effect of knocking down *ERG* on heptad genes in CD34+ HSPCs. *Right:* GSEA plots showing enrichment of genes associated with the Ery-precursor/Ery-precursor-like state in response to knocking down *ERG* in CD34^+^ HSPCs. FDR q-value for GSEA plots = 0 except where indicated by * q-value = 0.02.

Since the bulk of ME-1 cells were assigned as HSC-like, we reasoned that increased expression of *GATA2* might alter their trajectory towards the Ery-precursor-like state. We analysed RNAseq data from *GATA2* over-expression in ME-1 cells^61^ and found that increased *GATA2* led to increased *TAL1* and *RUNX1*, and reduced *ERG* and *LMO2*, similar to expression changes between Ery-precursor-like and HSC-like ME-1 cells(Figure 5C, *left*, compare to Figure 4F, *left*). GSEA analysis was used to compare GATA2 driven changes in global gene expression to expression differences between Ery-precursors and HSCs. Globally, genes that were high in Ery-precursors tended to go up following GATA2 overexpression, while genes that were low in Ery-precursors tended to go down (Figure 5C, *right*). *ERG* overexpression in CD34^+^ HSPCs promotes expansion of the progenitor pool^62^, and we have now shown that *ERG* expression is reduced across the HSC to Ery-precursor boundary in normal BM and ME-1 (Figure 4F). We therefore asked whether reduced *ERG* expression in CD34^+^ HSPCs promoted an Ery-progenitor phenotype. *ERG* knockdown led to downregulation of *FLI1*, *LYL1*, and *LMO2*, and upregulation of *GATA2* and *TAL1* (Figure 5D, *left*), similar to the expression changes observed for the HSC-Ery-progenitor transition in Palantir data (Figure 4F, *right*). GSEA analysis was used to compare *ERG* knockdown driven changes in global gene expression to expression differences between Ery-precursors and HSCs. Globally, genes that were high in Ery-precursors tended to go up following ERG knockdown, while genes that were low in Ery-precursors tended to go down (Figure 5D, *right*). Together, the perturbation data supports the notion that heptad genes, and in particular the triplet *GATA2*, *TAL1*, and *ERG*, form a functionally relevant interconnected network and play a key role in regulating cell state transitions in healthy blood and leukemic cells.

## DISCUSSION

Gene regulatory networks control cell fate decisions in development and disease. Here, we focused on heptad transcription factors and identified parallel phenotypes between healthy haematopoiesis and leukemic cells spanning single cell gene expression, chromatin state, and enhancer use (Figure 6A). Our data suggest that GATA2, TAL1, and ERG constitute a heptad sub-circuit that regulates stem cell to erythroid transition in healthy blood and leukemia (Figure 6B).

**Figure 6.**
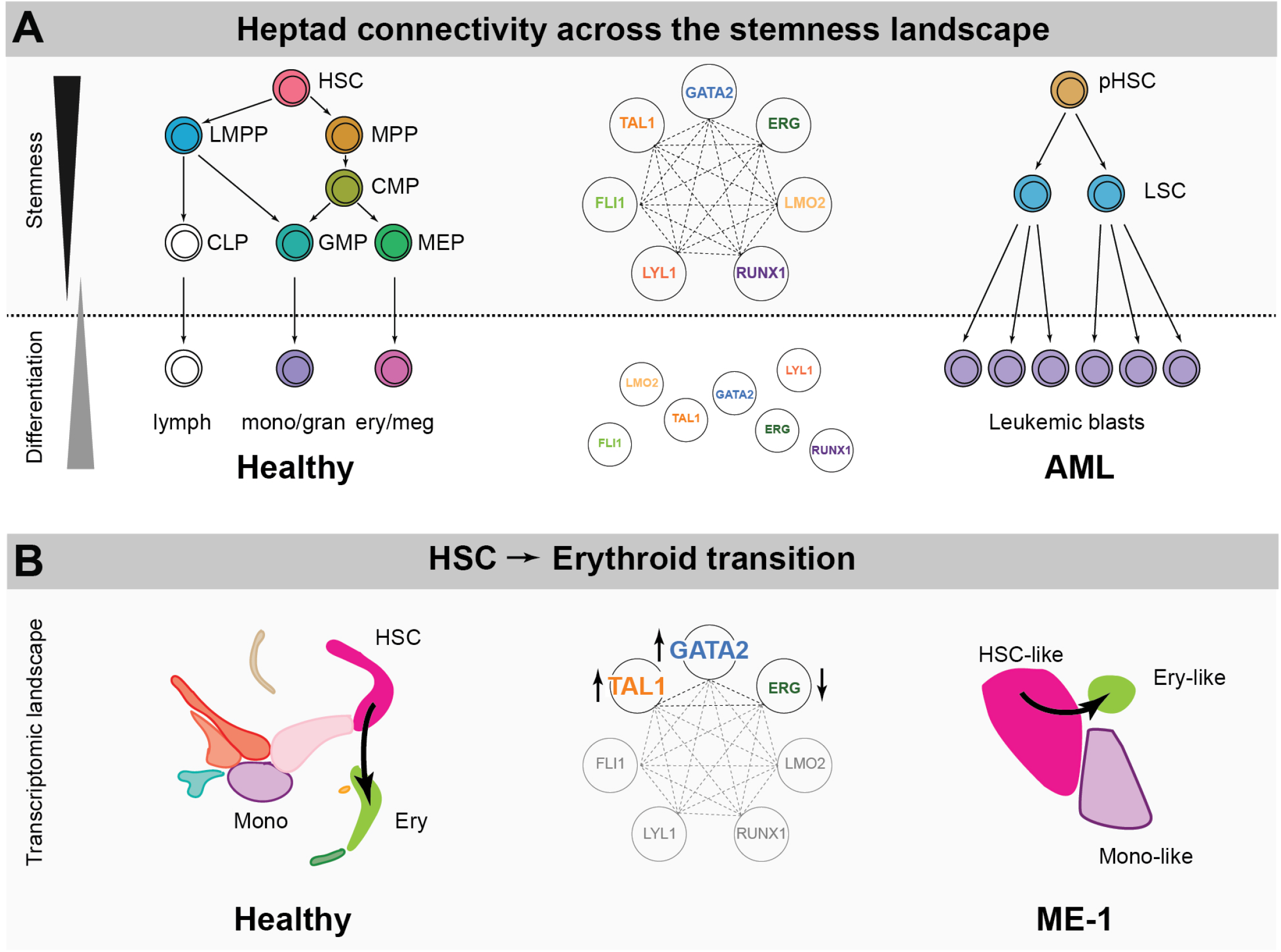
Proposed model of heptad activity across haematopoietic differentiation. (A) Heptad transcription factors form a densely interconnected network, with key regulatory elements accessible and heptad-bound in normal and leukemic stem cells. Accessibility of regulatory elements, and consequently heptad connectivity, is reduced as cells become more differentiated. (B) Schematics representing sc-RNAseq populations in normal and ME-1 cell populations. GATA2, TAL1, and ERG promote cell state changes along the HSC-Ery precursor axis in both normal CD34+ HSPCs and ME-1 cells.

### Insights into enhancer biology

Genome-wide chromatin state can be used to classify cell types^4^. Here we show that chromatin accessibility at only a handful of heptad enhancers could successfully classify all early stages of haematopoiesis as well as subpopulations of leukemic cells in AML. Most heptad enhancers were accessible in stem and progenitors and become selectively inaccessible at terminal differentiation, though exceptions were observed. In particular, we found that the *GATA2* −117 (mice: *Gata2* −77) enhancer was open only in CMPs and MEPs, suggesting a central role for this enhancer in erythroid transition and confirming previous murine models, where its deletion blocked erythroid and megakaryocytic differentiation^63^.

This enhancer has been previously studied in inv(3) AML where it is translocated close to oncogene *MECOM*/EVI1 leading to increased EVI1 and decreased *GATA2* expression^57,58^. We found that this enhancer was accessible in a subset of leukemic cells, and strongly heptad-bound in both AML cell lines compared to CD34+ HSPCs. In our reporter assays *GATA2* −117 also drove more luciferase reporter activity than *GATA2* +3.5, the other *GATA2* regulatory element. Thus, even in its normal genomic context *GATA2* −117 may play a role in driving *GATA2* expression in AML. Unlike *GATA2* −117, the *ERG* +85 enhancer was open in all HSPC subsets and across AML subtypes (Figure S1A). This enhancer has been linked to AML prognosis^43^ and has been used to identify LSCs within bulk AML populations^64,65^. Enhancers are replete with sequence motifs enabling binding of distinct TF families, either directly to DNA or indirectly via protein scaffolding, as observed for LMO2^66,67^ and RUNX1^42,44^. Here, we showed that evolutionarily conserved heptad enhancers rely heavily on ETS and GATA motifs, in agreement with previous reports that ETS-ETS-GATA motifs were enriched at blood enhancers^68^.

### Regulation of cell fate transitions by GATA2, TAL1 and ERG

Combinatorial binding of TFs is a key component of cell fate transitions^38^. We identify a triad of TFs-GATA2, TAL1, and ERG, whereby high *GATA2* and *TAL1*, and low *ERG* expression biased fate decisions towards the erythroid lineage in both CD34+ HSPCs and ME-1 leukemic cells. A similar circuit, comprised of *GATA2*, *TAL1*, and *FLI1* (an ETS TF closely related to ERG) has been previously reported during embryonic HSC specification^69^, while *GATA1*, *TAL1* and KLF1 form a sub-circuit in erythroid cells^70^. Indeed, recycling of regulatory modules is a key feature of developmental networks^38^, underlining the utility of cell classification strategies such as northstar^54^.

Each member of this triad is known to play complex roles in healthy blood and leukemia development. GATA2 levels control blood cell emergence in the embryonic aorta^71^, and is required for HSC maintenance^72^. Germline loss of function mutations in *GATA2* predisposes to MDS and AML^73^ and high *GATA2* expression is associated with poor prognosis in AML patients^74^. *TAL1* is also required for embryonic blood formation^48,75^ and drives erythroid and megakaryocytic differentiation programs^76^ but is not required for HSC maintenance^48,77,78^. However, dysregulation of TAL-1 is associated with T-ALL^48^. *ERG* is not required for HSC specification or differentiation but promotes HSC maintenance by restricting differentiation^79,80^. High *ERG* expression is a poor prognostic marker for AML^49,81–83^ and is leukemogenic in mouse models^84–87^, although its role in human leukemia is more subtle^62^.

### Clinical Implications

Therapeutic approaches to AML which force LSCs to differentiate have been sought^88^. Although TFs are relatively difficult drug targets, this might change as small molecules upregulating CEBPA^89,90^ or downregulating PU.1^91^ and RUNX1^92^ have been developed. Regulatory circuits such as the GATA2-TAL1-ERG triad described here may provide a conceptual framework within which to develop such therapies. A first approach would be to alter TF expression directly, as upregulating *GATA2* or downregulating *ERG* promotes erythroid differentiation. However, different leukemias may be primed towards specific differentiation pathways^93^. As such, *ERG* perturbation appears especially promising as this TF appears to preserve the progenitor state rather than bias towards a particular downstream fate. A second approach would be to focus on transcriptional regulators of these TFs. USP9X, a deubiquitinase that regulates ERG stability^94^ and is positively regulated by ERG in a feed forward loop is one such candidate^64^. A third approach would be to focus on specific enhancers such as *GATA2*-117, which is inaccessible in normal HSCs but open in the transitional progenitor states characteristic of AML, enabling preferential cytotoxicity in leukemic cells. Overall, a deeper understanding of heptad regulatory circuits in haematopoiesis and leukemia can help shape novel, data-based approaches to innovative cancer therapies.

## ACKNOWLEDGEMENTS

The authors thank the staff and donors of the Sydney Cord Blood Bank for providing cord bloods for research. Some of the data presented in this work was acquired by personnel and/or instruments at the Mark Wainwright Analytical Centre (MWAC) of UNSW Sydney, which is in part funded by the Research Infrastructure Programme of UNSW. The authors acknowledge the following funding support: JT was supported by the Anthony Rothe Memorial Trust; SS is supported by an International Postgraduate Student scholarship from UNSW and the Prince of Wales Clinical School; DB is supported by a Peter Doherty Fellowship from the National Health and Medical Research Council of Australia (APP1073768), a Cancer Institute NSW Early Career Fellowship, the Anthony Rothe Memorial Trust, and Gilead Sciences; BG is supported by a Wellcome Investigator award (206328/Z/17/Z); JEP is supported by the National Health and Medical Research Council of Australia (APP1042934, APP1102589, APP1008515), Translational Cancer Research Network - a Translational Cancer Research Centre funded by the Cancer Institute NSW, Anthony Rothe Memorial Trust, and philanthropic funding from Christina’s Light.

## AUTHORSHIP CONTRIBUTIONS

J.A.I.T., K.K., G.H., Y.H., J.S., D.R.C., S.S., and J.S. performed research and analysed data. D.C., A.S., D.B., and J.W.H.W. analysed data.

I.dJ. and J.L. provided key reagents.

B.G. and J.W.H.W. discussed and interpreted data.

J.A.I.T., F.Z., and J.P. conceived the study and wrote the paper.

## REFERENCES

1. Doulatov S, Notta F, Laurenti E, Dick JE. Hematopoiesis: a human perspective. Cell Stem Cell. 2012;10(2):120–136.

2. Velten L, Haas SF, Raffel S, et al. Human haematopoietic stem cell lineage commitment is a continuous process. Nat Cell Biol. 2017;19(4):271–281.

3. Buenrostro JD, Corces MR, Lareau CA, et al. Integrated Single-Cell Analysis Maps the Continuous Regulatory Landscape of Human Hematopoietic Differentiation. Cell. 2018;173(6):1535–1548 e1516.

4. Corces MR, Buenrostro JD, Wu B, et al. Lineage-specific and single-cell chromatin accessibility charts human hematopoiesis and leukemia evolution. Nat Genet. 2016;48(10):1193–1203.

5. Karamitros D, Stoilova B, Aboukhalil Z, et al. Single-cell analysis reveals the continuum of human lympho-myeloid progenitor cells. Nat Immunol. 2018;19(1):85–97.

6. Setty M, Kiseliovas V, Levine J, Gayoso A, Mazutis L, Pe’er D. Characterization of cell fate probabilities in single-cell data with Palantir. Nat Biotechnol. 2019;37(4):451–460.

7. Watcham S, Kucinski I, Gottgens B. New Insights into Haematopoietic Differentiation Landscapes from scRNA-seq. Blood. 2019.

8. Pimanda JE, Gottgens B. Gene regulatory networks governing haematopoietic stem cell development and identity. Int J Dev Biol. 2010;54(6-7):1201–1211.

9. Sive JI, Gottgens B. Transcriptional network control of normal and leukaemic haematopoiesis. Exp Cell Res. 2014;329(2):255–264.

10. Enver T, Pera M, Peterson C, Andrews PW. Stem cell states, fates, and the rules of attraction. Cell Stem Cell. 2009;4(5):387–397.

11. Moris N, Pina C, Arias AM. Transition states and cell fate decisions in epigenetic landscapes. Nat Rev Genet. 2016;17(11):693–703.

12. Wilkinson AC, Nakauchi H, Gottgens B. Mammalian Transcription Factor Networks: Recent Advances in Interrogating Biological Complexity. Cell Syst. 2017;5(4):319–331.

13. Thoms JAI, Beck D, Pimanda JE. Transcriptional networks in acute myeloid leukemia. Genes Chromosomes Cancer. 2019;58(12):859–874.

14. Dohner H, Weisdorf DJ, Bloomfield CD. Acute Myeloid Leukemia. N Engl J Med. 2015;373(12):1136–1152.

15. Horton SJ, Huntly BJ. Recent advances in acute myeloid leukemia stem cell biology. Haematologica. 2012;97(7):966–974.

16. Jan M, Snyder TM, Corces-Zimmerman MR, et al. Clonal evolution of preleukemic hematopoietic stem cells precedes human acute myeloid leukemia. Sci Transl Med. 2012;4(149):149ra118.

17. Shlush LI, Zandi S, Mitchell A, et al. Identification of pre-leukaemic haematopoietic stem cells in acute leukaemia. Nature. 2014;506(7488):328–333.

18. Basilico S, Gottgens B. Dysregulation of haematopoietic stem cell regulatory programs in acute myeloid leukaemia. J Mol Med (Berl). 2017;95(7):719–727.

19. Corces-Zimmerman MR, Hong WJ, Weissman IL, Medeiros BC, Majeti R. Preleukemic mutations in human acute myeloid leukemia affect epigenetic regulators and persist in remission. Proc Natl Acad Sci U S A. 2014;111(7):2548–2553.

20. Lapidot T, Sirard C, Vormoor J, et al. A cell initiating human acute myeloid leukaemia after transplantation into SCID mice. Nature. 1994;367(6464):645–648.

21. Goardon N, Marchi E, Atzberger A, et al. Coexistence of LMPP-like and GMP-like leukemia stem cells in acute myeloid leukemia. Cancer Cell. 2011;19(1):138–152.

22. Sarry JE, Murphy K, Perry R, et al. Human acute myelogenous leukemia stem cells are rare and heterogeneous when assayed in NOD/SCID/IL2Rgammac-deficient mice. J Clin Invest. 2011;121(1):384–395.

23. Shlush LI, Mitchell A, Heisler L, et al. Tracing the origins of relapse in acute myeloid leukaemia to stem cells. Nature. 2017;547(7661):104–108.

24. Eppert K, Takenaka K, Lechman ER, et al. Stem cell gene expression programs influence clinical outcome in human leukemia. Nat Med. 2011;17(9):1086–1093.

25. Gentles AJ, Plevritis SK, Majeti R, Alizadeh AA. Association of a leukemic stem cell gene expression signature with clinical outcomes in acute myeloid leukemia. JAMA. 2010;304(24):2706–2715.

26. Bonnet D, Dick JE. Human acute myeloid leukemia is organized as a hierarchy that originates from a primitive hematopoietic cell. Nat Med. 1997;3(7):730–737.

27. Sanz MA, Grimwade D, Tallman MS, et al. Management of acute promyelocytic leukemia: recommendations from an expert panel on behalf of the European LeukemiaNet. Blood. 2009;113(9):1875–1891.

28. DiNardo CD, Stein EM, de Botton S, et al. Durable Remissions with Ivosidenib in IDH1-Mutated Relapsed or Refractory AML. N Engl J Med. 2018;378(25):2386–2398.

29. Stein EM, DiNardo CD, Pollyea DA, et al. Enasidenib in mutant IDH2 relapsed or refractory acute myeloid leukemia. Blood. 2017;130(6):722–731.

30. Hansen E, Quivoron C, Straley K, et al. AG-120, an Oral, Selective, First-in-Class, Potent Inhibitor of Mutant IDH1, Reduces Intracellular 2HG and Induces Cellular Differentiation in TF-1 R132H Cells and Primary Human IDH1 Mutant AML Patient Samples Treated Ex Vivo. Blood. 2014;124(21):3734–3734.

31. Popovici-Muller J, Lemieux RM, Artin E, et al. Discovery of AG-120 (Ivosidenib): A First-in-Class Mutant IDH1 Inhibitor for the Treatment of IDH1 Mutant Cancers. ACS Medicinal Chemistry Letters. 2018;9(4):300–305.

32. Papaemmanuil E, Gerstung M, Bullinger L, et al. Genomic Classification and Prognosis in Acute Myeloid Leukemia. N Engl J Med. 2016;374(23):2209–2221.

33. Arber DA, Orazi A, Hasserjian R, et al. The 2016 revision to the World Health Organization classification of myeloid neoplasms and acute leukemia. Blood. 2016;127(20):2391–2405.

34. Cancer Genome Atlas Research N, Ley TJ, Miller C, et al. Genomic and epigenomic landscapes of adult de novo acute myeloid leukemia. N Engl J Med. 2013;368(22):2059–2074.

35. Assi SA, Imperato MR, Coleman DJL, et al. Subtype-specific regulatory network rewiring in acute myeloid leukemia. Nat Genet. 2019;51(1):151–162.

36. Yi G, Wierenga ATJ, Petraglia F, et al. Chromatin-Based Classification of Genetically Heterogeneous AMLs into Two Distinct Subtypes with Diverse Stemness Phenotypes. Cell Rep. 2019;26(4):1059–1069 e1056.

37. McKeown MR, Corces MR, Eaton ML, et al. Superenhancer Analysis Defines Novel Epigenomic Subtypes of Non-APL AML, Including an RARalpha Dependency Targetable by SY-1425, a Potent and Selective RARalpha Agonist. Cancer Discov. 2017;7(10):1136–1153.

38. Davidson EH. Emerging properties of animal gene regulatory networks. Nature. 2010;468(7326):911–920.

39. Bell CC, Fennell KA, Chan YC, et al. Targeting enhancer switching overcomes non-genetic drug resistance in acute myeloid leukaemia. Nat Commun. 2019;10(1):2723.

40. Fennell KA, Bell CC, Dawson MA. Epigenetic therapies in acute myeloid leukemia: where to from here? Blood. 2019;134(22):1891–1901.

41. Guo L, Li J, Zeng H, et al. A combination strategy targeting enhancer plasticity exerts synergistic lethality against BETi-resistant leukemia cells. Nat Commun. 2020;11(1):740.

42. Beck D, Thoms JA, Perera D, et al. Genome-wide analysis of transcriptional regulators in human HSPCs reveals a densely interconnected network of coding and noncoding genes. Blood. 2013;122(14):e12–22.

43. Diffner E, Beck D, Gudgin E, et al. Activity of a heptad of transcription factors is associated with stem cell programs and clinical outcome in acute myeloid leukemia. Blood. 2013;121(12):2289–2300.

44. Wilson NK, Foster SD, Wang X, et al. Combinatorial transcriptional control in blood stem/progenitor cells: genome-wide analysis of ten major transcriptional regulators. Cell Stem Cell. 2010;7(4):532–544.

45. Guibentif C, Ronn RE, Boiers C, et al. Single-Cell Analysis Identifies Distinct Stages of Human Endothelial-to-Hematopoietic Transition. Cell Rep. 2017;19(1):10–19.

46. Bergiers I, Andrews T, Vargel Bolukbasi O, et al. Single-cell transcriptomics reveals a new dynamical function of transcription factors during embryonic hematopoiesis. Elife. 2018;7.

47. Oram SH, Thoms JA, Pridans C, et al. A previously unrecognized promoter of LMO2 forms part of a transcriptional regulatory circuit mediating LMO2 expression in a subset of T-acute lymphoblastic leukaemia patients. Oncogene. 2010;29(43):5796–5808.

48. Curtis DJ, Salmon JM, Pimanda JE. Concise review: Blood relatives: formation and regulation of hematopoietic stem cells by the basic helix-loop-helix transcription factors stem cell leukemia and lymphoblastic leukemia-derived sequence 1. Stem Cells. 2012;30(6):1053–1058.

49. Marcucci G, Baldus CD, Ruppert AS, et al. Overexpression of the ETS-related gene, ERG, predicts a worse outcome in acute myeloid leukemia with normal karyotype: a Cancer and Leukemia Group B study. J Clin Oncol. 2005;23(36):9234–9242.

50. Li Y, Luo H, Liu T, Zacksenhaus E, Ben-David Y. The ets transcription factor Fli-1 in development, cancer and disease. Oncogene. 2015;34(16):2022–2031.

51. Mandoli A, Singh AA, Jansen PW, et al. CBFB-MYH11/RUNX1 together with a compendium of hematopoietic regulators, chromatin modifiers and basal transcription factors occupies self-renewal genes in inv(16) acute myeloid leukemia. Leukemia. 2014;28(4):770–778.

52. Mandoli A, Singh AA, Prange KHM, et al. The Hematopoietic Transcription Factors RUNX1 and ERG Prevent AML1-ETO Oncogene Overexpression and Onset of the Apoptosis Program in t(8;21) AMLs. Cell Rep. 2016;17(8):2087–2100.

53. Sotoca AM, Prange KH, Reijnders B, et al. The oncofusion protein FUS-ERG targets key hematopoietic regulators and modulates the all-trans retinoic acid signaling pathway in t(16;21) acute myeloid leukemia. Oncogene. 2016;35(15):1965–1976.

54. Zanini F, Berghuis BA, Jones RC, et al. Northstar enables automatic classification of known and novel cell types from tumor samples. Sci Rep. 2020;10(1):15251.

55. La Manno G, Soldatov R, Zeisel A, et al. RNA velocity of single cells. Nature. 2018;560(7719):494–498.

56. Bergen V, Lange M, Peidli S, Wolf FA, Theis FJ. Generalizing RNA velocity to transient cell states through dynamical modeling. Nat Biotechnol. 2020.

57. Yamazaki H, Suzuki M, Otsuki A, et al. A remote GATA2 hematopoietic enhancer drives leukemogenesis in inv(3)(q21;q26) by activating EVI1 expression. Cancer Cell. 2014;25(4):415–427.

58. Groschel S, Sanders MA, Hoogenboezem R, et al. A single oncogenic enhancer rearrangement causes concomitant EVI1 and GATA2 deregulation in leukemia. Cell. 2014;157(2):369–381.

59. Rucker FG, Sander S, Dohner K, Dohner H, Pollack JR, Bullinger L. Molecular profiling reveals myeloid leukemia cell lines to be faithful model systems characterized by distinct genomic aberrations. Leukemia. 2006;20(6):994–1001.

60. Yanagisawa K, Horiuchi T, Fujita S. Establishment and characterization of a new human leukemia cell line derived from M4E0. Blood. 1991;78(2):451–457.

61. Yi G, Mandoli A, Jussen L, et al. CBFbeta-MYH11 interferes with megakaryocyte differentiation via modulating a gene program that includes GATA2 and KLF1. Blood Cancer J. 2019;9(3):33.

62. Tursky ML, Beck D, Thoms JA, et al. Overexpression of ERG in cord blood progenitors promotes expansion and recapitulates molecular signatures of high ERG leukemias. Leukemia. 2015;29(4):819–827.

63. Johnson KD, Conn DJ, Shishkova E, et al. Constructing and deconstructing GATA2-regulated cell fate programs to establish developmental trajectories. J Exp Med. 2020;217(11).

64. Aqaqe N, Yassin M, Yassin AA, et al. An ERG Enhancer-Based Reporter Identifies Leukemia Cells with Elevated Leukemogenic Potential Driven by ERG-USP9X Feed-Forward Regulation. Cancer Res. 2019;79(15):3862–3876.

65. Yassin M, Aqaqe N, Yassin AA, et al. A novel method for detecting the cellular stemness state in normal and leukemic human hematopoietic cells can predict disease outcome and drug sensitivity. Leukemia. 2019;33(8):2061–2077.

66. Osada H, Grutz G, Axelson H, Forster A, Rabbitts TH. Association of erythroid transcription factors: complexes involving the LIM protein RBTN2 and the zinc-finger protein GATA1. Proc Natl Acad Sci U S A. 1995;92(21):9585–9589.

67. Wadman I, Li J, Bash RO, et al. Specific in vivo association between the bHLH and LIM proteins implicated in human T cell leukemia. EMBO J. 1994;13(20):4831–4839.

68. Donaldson IJ, Chapman M, Kinston S, et al. Genome-wide identification of cis-regulatory sequences controlling blood and endothelial development. Hum Mol Genet. 2005;14(5):595–601.

69. Pimanda JE, Ottersbach K, Knezevic K, et al. Gata2, Fli1, and Scl form a recursively wired gene-regulatory circuit during early hematopoietic development. Proc Natl Acad Sci U S A. 2007;104(45):17692–17697.

70. Wontakal SN, Guo X, Smith C, et al. A core erythroid transcriptional network is repressed by a master regulator of myelo-lymphoid differentiation. Proc Natl Acad Sci U S A. 2012;109(10):3832–3837.

71. Eich C, Arlt J, Vink CS, et al. In vivo single cell analysis reveals Gata2 dynamics in cells transitioning to hematopoietic fate. J Exp Med. 2018;215(1):233–248.

72. Menendez-Gonzalez JB, Vukovic M, Abdelfattah A, et al. Gata2 as a Crucial Regulator of Stem Cells in Adult Hematopoiesis and Acute Myeloid Leukemia. Stem Cell Reports. 2019;13(2):291–306.

73. Hahn CN, Chong CE, Carmichael CL, et al. Heritable GATA2 mutations associated with familial myelodysplastic syndrome and acute myeloid leukemia. Nat Genet. 2011;43(10):1012–1017.

74. Vicente C, Vazquez I, Conchillo A, et al. Overexpression of GATA2 predicts an adverse prognosis for patients with acute myeloid leukemia and it is associated with distinct molecular abnormalities. Leukemia. 2012;26(3):550–554.

75. Lancrin C, Sroczynska P, Stephenson C, Allen T, Kouskoff V, Lacaud G. The haemangioblast generates haematopoietic cells through a haemogenic endothelium stage. Nature. 2009;457(7231):892–895.

76. Elwood NJ, Zogos H, Pereira DS, Dick JE, Begley CG. Enhanced megakaryocyte and erythroid development from normal human CD34(+) cells: consequence of enforced expression of SCL. Blood. 1998;91(10):3756–3765.

77. Mikkola HK, Klintman J, Yang H, et al. Haematopoietic stem cells retain long-term repopulating activity and multipotency in the absence of stem-cell leukaemia SCL/tal-1 gene. Nature. 2003;421(6922):547–551.

78. Robertson SM, Kennedy M, Shannon JM, Keller G. A transitional stage in the commitment of mesoderm to hematopoiesis requiring the transcription factor SCL/tal-1. Development. 2000;127(11):2447–2459.

79. Taoudi S, Bee T, Hilton A, et al. ERG dependence distinguishes developmental control of hematopoietic stem cell maintenance from hematopoietic specification. Genes Dev. 2011;25(3):251–262.

80. Knudsen KJ, Rehn M, Hasemann MS, et al. ERG promotes the maintenance of hematopoietic stem cells by restricting their differentiation. Genes Dev. 2015;29(18):1915–1929.

81. Marcucci G, Maharry K, Whitman SP, et al. High expression levels of the ETS-related gene, ERG, predict adverse outcome and improve molecular risk-based classification of cytogenetically normal acute myeloid leukemia: a Cancer and Leukemia Group B Study. J Clin Oncol. 2007;25(22):3337–3343.

82. Schwind S, Marcucci G, Maharry K, et al. BAALC and ERG expression levels are associated with outcome and distinct gene and microRNA expression profiles in older patients with de novo cytogenetically normal acute myeloid leukemia: a Cancer and Leukemia Group B study. Blood. 2010;116(25):5660–5669.

83. Metzeler KH, Dufour A, Benthaus T, et al. ERG expression is an independent prognostic factor and allows refined risk stratification in cytogenetically normal acute myeloid leukemia: a comprehensive analysis of ERG, MN1, and BAALC transcript levels using oligonucleotide microarrays. J Clin Oncol. 2009;27(30):5031–5038.

84. Thoms JA, Birger Y, Foster S, et al. ERG promotes T-acute lymphoblastic leukemia and is transcriptionally regulated in leukemic cells by a stem cell enhancer. Blood. 2011;117(26):7079–7089.

85. Goldberg L, Tijssen MR, Birger Y, et al. Genome-scale expression and transcription factor binding profiles reveal therapeutic targets in transgenic ERG myeloid leukemia. Blood. 2013;122(15):2694–2703.

86. Carmichael CL, Metcalf D, Henley KJ, et al. Hematopoietic overexpression of the transcription factor Erg induces lymphoid and erythro-megakaryocytic leukemia. Proc Natl Acad Sci U S A. 2012;109(38):15437–15442.

87. Salek-Ardakani S, Smooha G, de Boer J, et al. ERG is a megakaryocytic oncogene. Cancer Res. 2009;69(11):4665–4673.

88. Nowak D, Stewart D, Koeffler HP. Differentiation therapy of leukemia: 3 decades of development. Blood. 2009;113(16):3655–3665.

89. Namasu CY, Katzerke C, Brauer-Hartmann D, et al. ABR, a novel inducer of transcription factor C/EBPalpha, contributes to myeloid differentiation and is a favorable prognostic factor in acute myeloid leukemia. Oncotarget. 2017;8(61):103626–103639.

90. Radomska HS, Jernigan F, Nakayama S, et al. A Cell-Based High-Throughput Screening for Inducers of Myeloid Differentiation. J Biomol Screen. 2015;20(9):1150–1159.

91. Antony-Debre I, Paul A, Leite J, et al. Pharmacological inhibition of the transcription factor PU.1 in leukemia. J Clin Invest. 2017;127(12):4297–4313.

92. Morita K, Suzuki K, Maeda S, et al. Genetic regulation of the RUNX transcription factor family has antitumor effects. J Clin Invest. 2017;127(7):2815–2828.

93. van Galen P, Hovestadt V, Wadsworth Ii MH, et al. Single-Cell RNA-Seq Reveals AML Hierarchies Relevant to Disease Progression and Immunity. Cell. 2019;176(6):1265–1281 e1224.

94. Wang S, Kollipara RK, Srivastava N, et al. Ablation of the oncogenic transcription factor ERG by deubiquitinase inhibition in prostate cancer. Proc Natl Acad Sci U S A. 2014;111(11):4251–4256.

