## Supplementary methods and figures for "Disruption of a GATA2, TAL1, ERG regulatory circuit promotes erythroid transition in healthy and leukemic stem cells"

#### Cell Culture, Transfection, and Lentiviral Transduction

ME-1 and KG-1 cells were from DSMZ (Germany). Stocks were STR confirmed in 2015 and tested for mycoplasma monthly. Cells were grown in RPMI1640 supplemented with 20% (ME-1) or 10% (KG-1) fetal bovine serum, 2mM GlutaMAX, and pen/strep (all Life Technologies). Cord bloods were obtained from Sydney Cord Blood Bank under ethics approval 08/190 from South Eastern Sydney Local Health District, and $CD34^+$ cells isolated using MACS beads[1] (Miltenyi Biotec). Experiments using CD34+ cells were performed on pooled cells from multiple donors.

Transient transfections were performed, and luciferase activity measured and normalised to β-galactosidase activity as previously described[2]. Lentiviral shRNA vectors (shLUC cat# SHC007, shERG cat# SHCLND clone TRCN0000431367) were modified to contain a GFP reporter instead of a puromycin resistance cassette, and lentiviral production and transduction performed essentially as described [3]. GFP+ transduced cells were sorted for RNA extraction 72-96 hours following transduction, and RNA extracted using miRNeasy mini kit (Qiagen).

#### Luciferase Reporter Assays

Details of regulatory regions cloned are in Table S4. Corresponding dog (Cf), mouse (Mm) and opossum (Md) sequences were identified using blat (UCSC browser) and aligned using Geneious Prime (https://www.geneious.com). TF consensus motifs that were conserved across at least 3/4 species (or 2/3 where conservation did not extend to 4 species) were considered conserved and selected for mutational analysis. Conserved TF binding consensus motifs were generally mutated as follows; CANNTG to CTCGAG, GGAW to TCGA, GATA to TCGA, and TGYGGT to CTCGAG. Mutants were checked to ensure mutations did not introduce a new motif. Regions were either synthesised (Thermo Fisher) or PCR amplified and mutated using site directed mutagenesis then cloned into pGL2 promoter or basic vectors (Promega).

At least two independent experiments with triplicate measurements, were performed for each construct. Luciferase was normalised to β-galactosidase then scaled to mean value of the empty vector for that experiment = 1. Statistical comparisons used a Holm-Sidak t-test with alpha = 0.05, * indicates $P < 0.05$, ** indicates $P < 0.01$, *** indicates $P < 0.001$ (GraphPad Prism). For composite heatmaps, luciferase readings were normalised to β-galactosidase, then scaled such that the mean value of the WT replicates was 100. Data from all experiments was then averaged and scaled to mean value of the empty vector = 1.

**Quantitative RT-PCR**

RNA was reverse transcribed using M-MLV reverse transcriptase (Thermo Fisher) and quantitative PCR using PowerUp SYBR Green Master Mix (Thermo Fisher) and a MX3000P thermocycler (Stratagene). Primer sequences are listed in Table S7. Seven reference genes (β-actin, B2M, HPRT, SDHA, YWHAZ, EMC7, PSMB4[4,5]) were screened for level of variance across all samples using qbase+ software (Biogazelle, Belgium). Relative expression of heptad genes was quantified using qbase+ software (Biogazelle, Belgium) with β-actin, EMC7, and PSMB4 as reference genes. Data was graphed in Python 3.7 using the Matplotlib[6] and seaborn (https://github.com/mwaskom/seaborn/tree/v0.8.1) modules.

**ATACseq, DHSseq, and ChIPseq data processing**

ATACseq fastq files (GSE74912[7]; Table S3) were downloaded and processed as follows; remove adaptors (cutadapt), align to the hg38 human genome build (bwa mem), remove mitochondrial reads (grep chrM), remove duplicate reads (picard MarkDuplicates using options REMOVE_DUPLICATES=TRUE REMOVE_SEQUENCING_DUPLICATES=TRUE), shift alignment for Tn5 (alignmentSieve using options --minMappingQuality 10 --ATACshift), and create bigwig files (bamCoverage using options --binSize 10 –exactScaling).

Wiggle files corresponding to DHS data aligned to the hg38 human genome build (GSE108316[8]; Table S3) were downloaded from GEO and converted to bigwig format using wigToBigWig[9].

Public (GSM1816075[10], GSE45144[11], GSE46044[12]; Table S3) and our ChIPseq fastq files were processed as follows; remove adaptors (cutadapt), align to the hg38 human genome build (bwa mem), remove duplicate reads (picard MarkDuplicates using options REMOVE_DUPLICATES=TRUE), and create bigwig files (bamCoverage using options --binSize 10 --exactScaling (--extendReads 200 for SE data only)).

**ChIPseq peak calling**

Global ChIP-seq peaks were called against the negative control IgG using three publicly available peak finding programs, MACS2[13] (p-value 0.00001), findPeaks (HOMER[14], FDR 0.05), and SPP[15] (FDR 0.05). Peaks called by at least two algorithms were identified using the BEDTools package (v2.27.1)[16] as high confidence binding sites for downstream analysis. To determine the presence of ChIP-seq peaks at selected gene regulatory elements, we used BEDTools to generate a binary data matrix by first merging the coordinates of all input files (bedtools merge), followed by applying the intersectBed -c command to determine overlap. Regulatory network connectivity was inferred from called peaks, and represented using BioTapestry software[17].

**Single cell RNA sequencing**

Single cell RNA-seq libraries were prepared with approximately 2000 cells using the Chromium Single Cell 3' Library and Gel Bead Kit v2 (10X Genomics) as per manufacturer's protocol. cDNA amplification was undertaken for 11 cycles and the subsequent sample index (SI) PCR for 12 cycles. Libraries were sequenced using the Illumina NextSeq500 platform with 150bp PE reads (Illumina) at a depth of approximately 100K reads/cell. Reads were aligned to the human reference genome (GRCh38) and unique molecular identifier (UMI) count matrices generated using CellRanger software version 2.0.1 (10X Genomics).

**scRNAseq Analysis**

The whole secondary analysis for Figs 1 and 4 is included at https://github.com/iosonofabio/heptad_paper. Single cell RNA-Seq data on healthy hematopoietic cells was downloaded from the Palantir github repository as described on https://github.com/dpeerlab/Palantir/blob/master/README.md, Rep1. Embedding coordinates, colors, cluster metadata, and smoothed counts data were extracted from the h5ad file and plotted using singlet (https://github.com/iosonofabio/singlet).

Count and metadata tables from CellRanger (10X Genomics) were converted to loom format (http://loompy.org/). and normalised to "counts per ten thousand (uniquely mapped) reads" as customary for droplet-based scRNA-Seq data and cumulative distributions plotted from this normalised data. Spearman correlations between gene expressions were computed via singlet (see above) using scipy's "average" (i.e. default) tie breaker for dropouts. The symmetric correlation matrix was ordered by hierarchical (average linkage) clustering on L2 distance with additional optimal leaf ordering and plotted as a heatmap. Conditional distributions of gene expression were computed via quantiles followed by Gaussian kernel density estimate in logarithmic space with a kernel bandwidth determined by the Scott method (scipy's default).

Palantir data were loaded as described above, normalised using the same approach as our ME1 data, and subsampled to 40 cells per cell type using singlet. northstar's subsample method[18] was used to infer cell states within ME-1 guided by the Palantir data[19]. In the graph construction step, 10 external (non-mutual) neighbors were allowed in order to compensate for the fact that ME1 cells are, on an absolute scale, quite distant from actual hematopoietic cells. ME1 cells were then colored based on northstar's community assignments to match their colors in Fig 1 (and the Palantir paper). RNA velocity[20] was computed using scVelo[21] and projected onto northstar's hybrid embedding. Gene expression was plotted in the same embedding after iterative nearest-neighbor smoothing, similar to the treatment by the original Palantir authors on their data, to enhance patterns in a context of overall low expression. For the prediction of ME1 cell state based on heptad and subsets thereof, we trained a random forest classifier using scikit-learn and evaluated its performance via train/test splits. Sample balancing and forest parameters were explored and yielded overall similar results – with the expected trade-offs in terms of specificity/sensitivity. Bar plots were used to represent the results.

**RNA sequencing and analysis**

Fastq files were aligned to the hg19 human reference genome using TopHat2 with standard parameters[22] and differential gene expression determined using DeSeq2 with standard parameters and paired sample information included as a regression variable[23].

**Gene Set Enrichment Analysis (GSEA)**

Gene Set Enrichment Analysis was performed using the GSEAPreranked module of GSEA software version 4.0.3[24] (Broad Institute). Differential gene expression lists for GATA2 overexpression in ME1 and ERG knockdown in CD34+ were filtered for genes that appeared in both lists and ranked by log2 fold change. Genelists for genes UP or DOWN in Ery-like compared to HSC-like were derived from differential expression analysis of scRNAseq datasets (ME-1 or Palantir, Table S8).

**LIST OF SUPPLEMENTARY TABLES**

Table S1 – Antibodies used for ChIP

Table S2 – hg38 chromosomal coordinates for heptad regulatory regions

Table S3 – Previously published datasets used in this manuscript

Table S4 – Sequences of normal and mutant heptad regulatory regions used for luciferase assays

Table S5 – Expression changes between HSC-like and Ery-precursor-like ME-1 cells

Table S6 – Expression changes between HSC and Ery-precursor healthy BM cells

Table S7 – primers used for PCR

Table S8 – GSEA genelists

**SUPPLEMENTARY FIGURES**

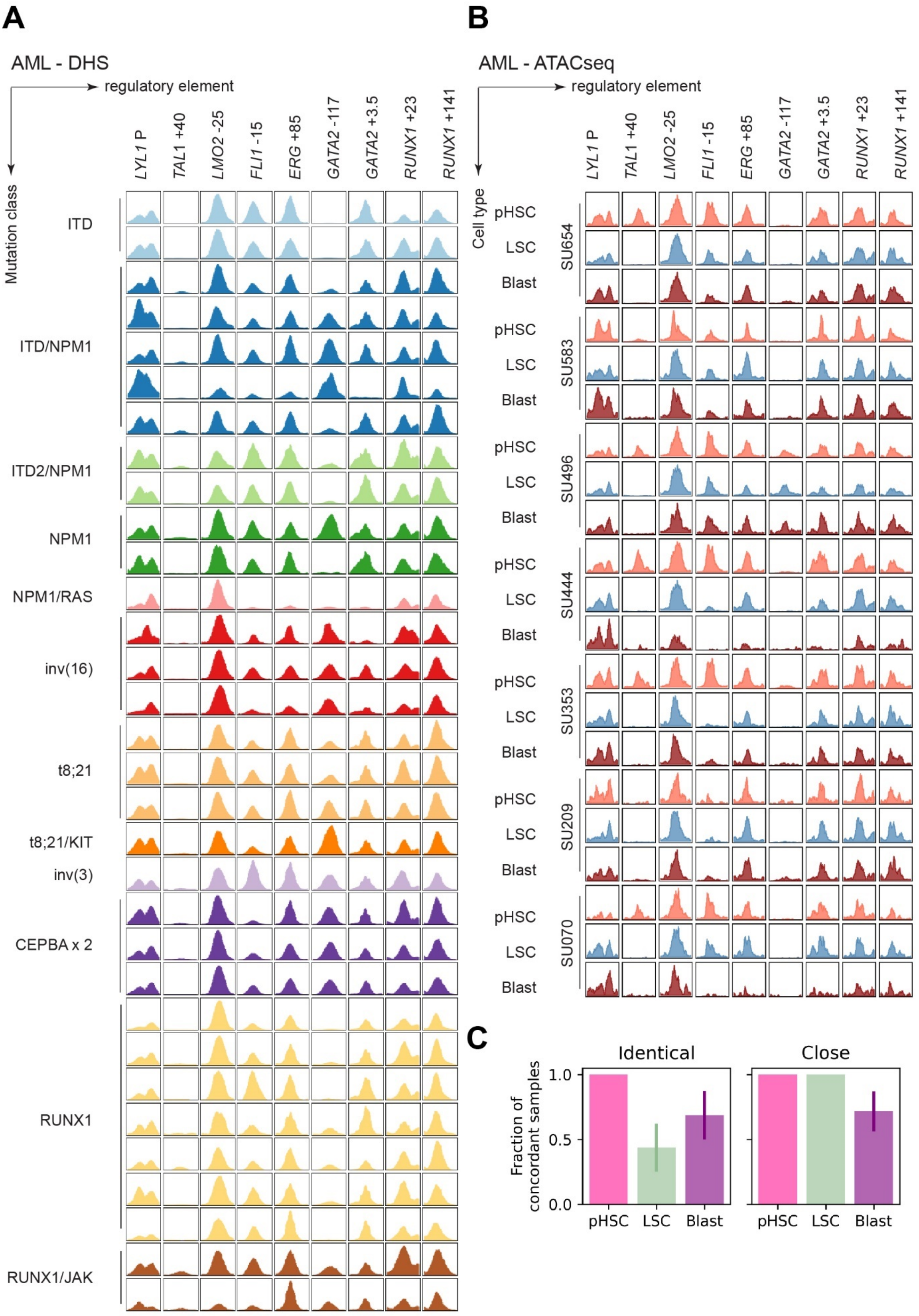
A
AML - DHS
regulatory element
Mutation class
LYL1 P
TAL1 +40
LMO2 -25
FLI1 -15
ERG +85
GATA2 -117
GATA2 +3.5
RUNX1 +23
RUNX1 +141
ITD
ITD/NPM1
ITD2/NPM1
NPM1
NPM1/RAS
inv(16)
t8;21
t8;21/KIT
inv(3)
CEPBA x 2
RUNX1
RUNX1/JAK
B
AML - ATACseq
regulatory element
Cell type
pHSC
LSC
Blast
SU654
SU583
SU496
SU444
SU353
SU209
SU070
C
Identical
Close
Fraction of concordant samples
1.0
0.5
0.0
pHSC
LSC
Blast

Figure S1

**Supplementary Figure 1 – Accessibility profiles in AML cells**

(A) DHS profiles at heptad regulatory regions for a range of AML mutational subtypes. Each row is a separate sample, samples are coloured by mutational subtype. (B) ATACseq profiles at heptad regulatory regions of seven AML patients, showing pre-leukemic HSCs (pHSC), leukemic stem cells (LSC), and leukemic blasts (Blast). (C) Fraction of samples assigned to the same cell type in our heptad classifier as in the genome-wide enhancer scan[7] as in Figure 1H. 100 bootstraps over patients were performed and the mean and standard deviations are shown. In the right chart, GMP and LMPP are counted as a single cell type due to their similar heptad enhancer profile.

# CD34

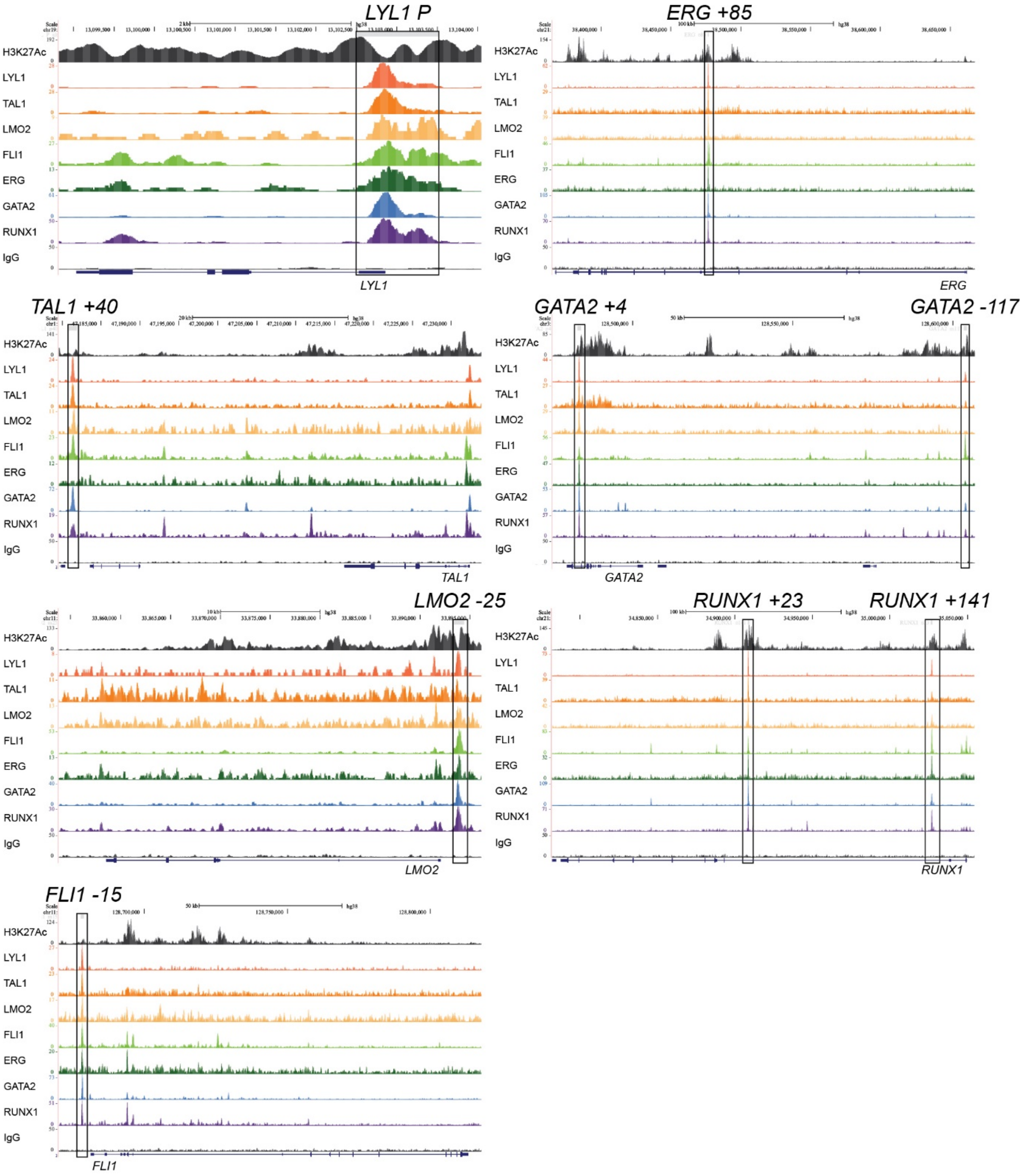


Figure S2

**Supplementary Figure 2 – Locus-wide ChIPseq profiles in CD34$^+$ cells**

ChIPseq profiles of H3K27Ac[10], heptad transcription factors, and IgG at heptad gene loci in CD34$^+$ HSPCs. Heptad regulatory elements are marked with black boxes. The genomic regions (hg38) plotted are as follows; TAL1 - chr1:47,179,588-47,233,861, LMO2 - chr11:33,853,840-33,896,143, FLI1 - chr11:128,670,163-128,817,804, LYL1 - chr19:13,098,820-13,104,046, RUNX1 - chr21:34,782,153-35,054,418, ERG - chr21:38,365,284-38,667,777, GATA2 - chr3:128,475,026-128,606,928.

# ME-1

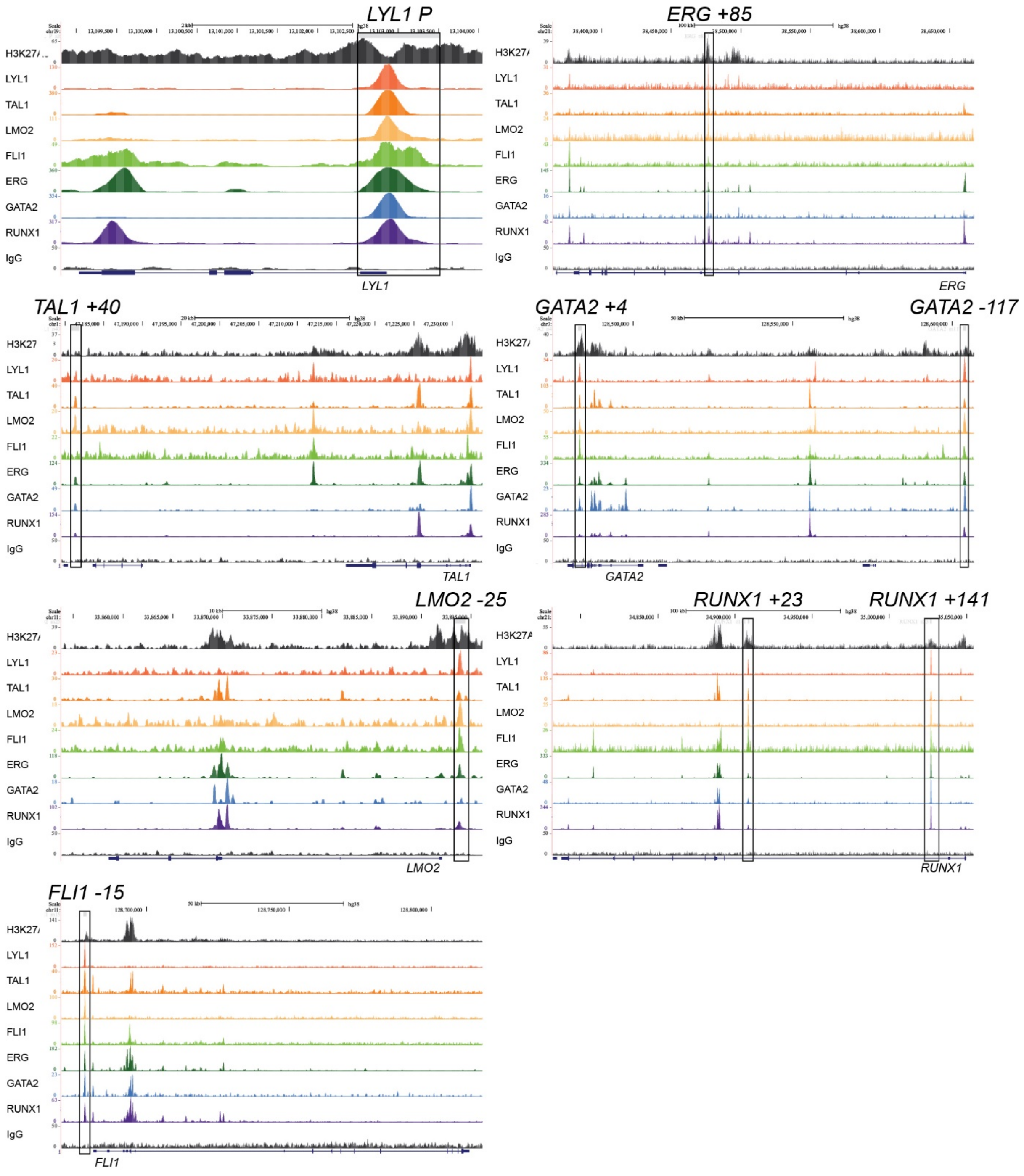


Figure S3

**Supplementary Figure 3 – Locus-wide ChIPseq profiles in ME-1 cells**

ChIPseq profiles of H3K27Ac, heptad transcription factors, and IgG at heptad gene loci in ME-1 cells. Heptad regulatory elements are marked with black boxes. The genomic regions (hg38) plotted are as follows; TAL1 - chr1:47,179,588-47,233,861, LMO2 - chr11:33,853,840-33,896,143, FLI1 - chr11:128,670,163-128,817,804, LYL1 - chr19:13,098,820-13,104,046, RUNX1 - chr21:34,782,153-35,054,418, ERG - chr21:38,365,284-38,667,777, GATA2 - chr3:128,475,026-128,606,928.

# KG-1

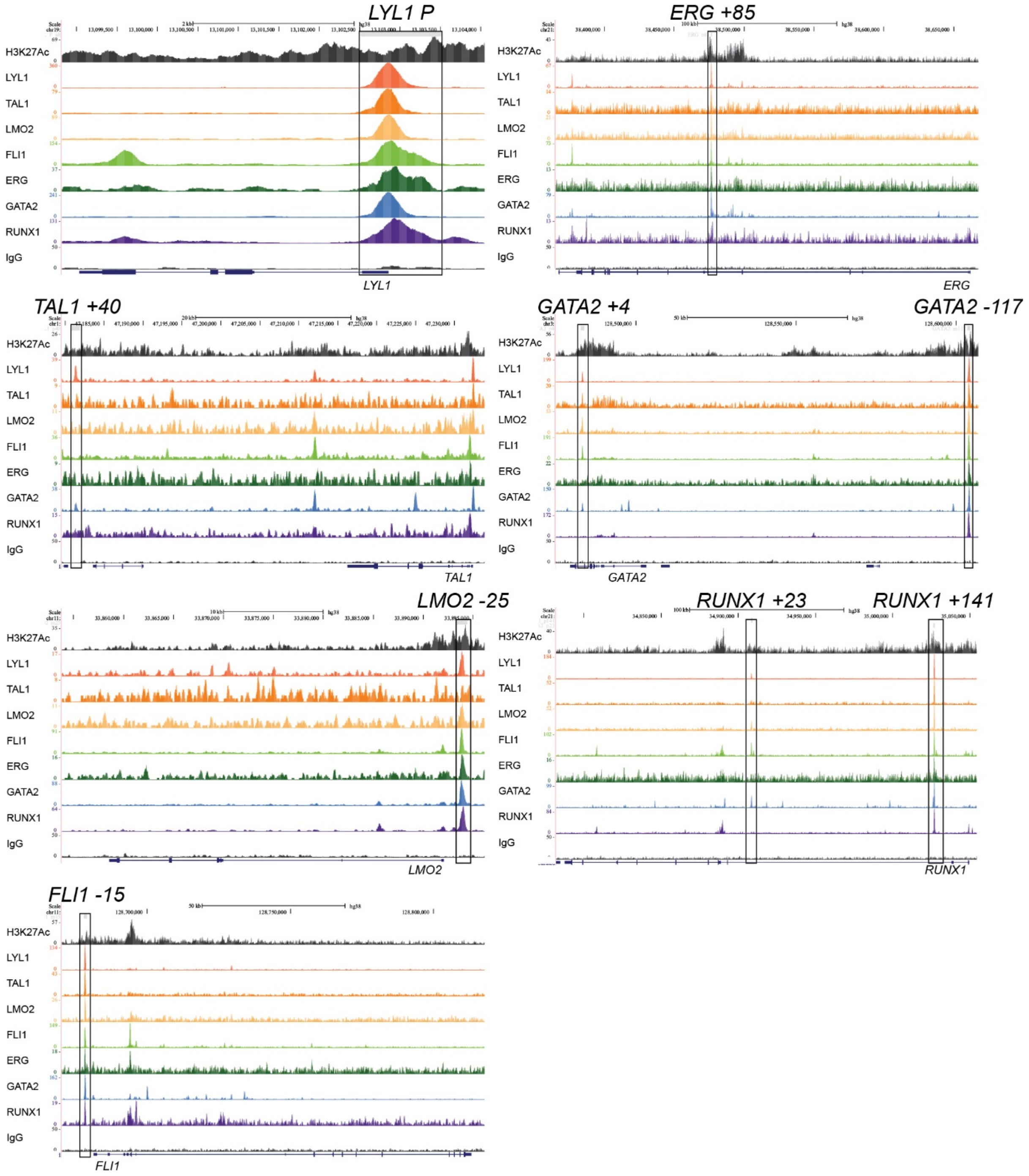


Figure S4

**Supplementary Figure 4 – Locus-wide ChIPseq profiles in KG-1 cells**

ChIPseq profiles of H3K27Ac, heptad transcription factors, and IgG at heptad gene loci in KG-1 cells. Heptad regulatory elements are marked with black boxes. The genomic regions (hg38) plotted are as follows; TAL1 - chr1:47,179,588-47,233,861, LMO2 - chr11:33,853,840-33,896,143, FLI1 - chr11:128,670,163-128,817,804, LYL1 - chr19:13,098,820-13,104,046, RUNX1 - chr21:34,782,153-35,054,418, ERG - chr21:38,365,284-38,667,777, GATA2 - chr3:128,475,026-128,606,928.

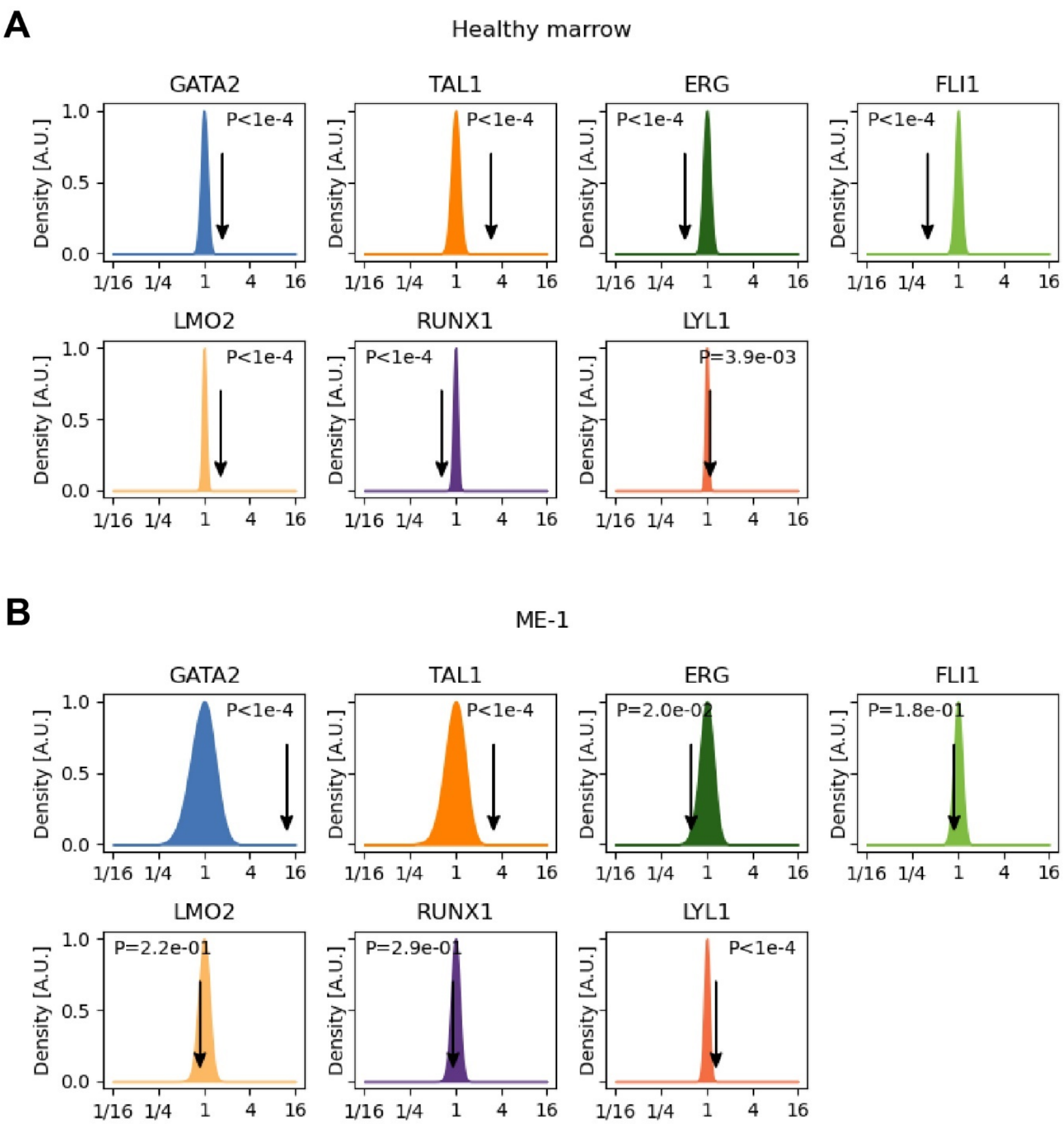
A
Healthy marrow
GATA2
TAL1
ERG
FLI1
LMO2
RUNX1
LYL1
Density [A.U.]
P<1e-4
P=3.9e-03
1/16 1/4 1 4 16
B
ME-1
GATA2
TAL1
ERG
FLI1
LMO2
RUNX1
LYL1
Density [A.U.]
P<1e-4
P=2.0e-02
P=1.8e-01
P=2.2e-01
P=2.9e-01
1/16 1/4 1 4 16

Figure S5

**Supplementary Figure 5 – Expression differences between HSC-like and Ery precursor-like cells in normal BM and ME-1 cells**

Distribution of heptad gene expression ratios between HSC-like and Ery precursor-like over 10,000 randomisations of cell state identity. P values indicate what fraction of randomisations have more extreme ratios than those observed in real data (one-tailed). P<1e-4 indicates that not a single simulation had ratios as imbalanced as the data. (A) Normal BM. All TFs are significant, with *LYL1* having the smallest effect size and largest P value. (B) ME-1 cells. *GATA2, ERG, TAL1*, and *LYL1* are significant below 0.05, with *LYL1* having the smallest effect size.

### REFERENCES


1. Tursky ML, Beck D, Thoms JA, et al. Overexpression of ERG in cord blood progenitors promotes expansion and recapitulates molecular signatures of high ERG leukemias. *Leukemia*. 2015;29(4):819-827.
2. Diffner E, Beck D, Gudgin E, et al. Activity of a heptad of transcription factors is associated with stem cell programs and clinical outcome in acute myeloid leukemia. *Blood*. 2013;121(12):2289-2300.
3. Unnikrishnan A, Guan YF, Huang Y, et al. A quantitative proteomics approach identifies ETV6 and IKZF1 as new regulators of an ERG-driven transcriptional network. *Nucleic Acids Res*. 2016;44(22):10644-10661.
4. Vandesompele J, De Preter K, Pattyn F, et al. Accurate normalization of real-time quantitative RT-PCR data by geometric averaging of multiple internal control genes. *Genome Biol*. 2002;3(7):RESEARCH0034.
5. Eisenberg E, Levanon EY. Human housekeeping genes, revisited. *Trends Genet*. 2013;29(10):569-574.
6. Hunter JD. Matplotlib is a 2D graphics package used for Python for application development, interactive scripting, and publication-quality image generation across user interfaces and operating systems. *Computing in Science & Engineering*. 2007;9:90--95.
7. Corces MR, Buenrostro JD, Wu B, et al. Lineage-specific and single-cell chromatin accessibility charts human hematopoiesis and leukemia evolution. *Nat Genet*. 2016;48(10):1193-1203.
8. Assi SA, Imperato MR, Coleman DJL, et al. Subtype-specific regulatory network rewiring in acute myeloid leukemia. *Nat Genet*. 2019;51(1):151-162.
9. Kent WJ, Zweig AS, Barber G, Hinrichs AS, Karolchik D. BigWig and BigBed: enabling browsing of large distributed datasets. *Bioinformatics*. 2010;26(17):2204-2207.
10. Huang J, Liu X, Li D, et al. Dynamic Control of Enhancer Repertoires Drives Lineage and Stage-Specific Transcription during Hematopoiesis. *Dev Cell*. 2016;36(1):9-23.
11. Beck D, Thoms JA, Perera D, et al. Genome-wide analysis of transcriptional regulators in human HSPCs reveals a densely interconnected network of coding and noncoding genes. *Blood*. 2013;122(14):e12-22.
12. Mandoli A, Singh AA, Jansen PW, et al. CBFB-MYH11/RUNX1 together with a compendium of hematopoietic regulators, chromatin modifiers and basal transcription factors occupies self-renewal genes in inv(16) acute myeloid leukemia. *Leukemia*. 2014;28(4):770-778.
13. Zhang Y, Liu T, Meyer CA, et al. Model-based analysis of ChIP-Seq (MACS). *Genome Biol*. 2008;9(9):R137.
14. Heinz S, Benner C, Spann N, et al. Simple combinations of lineage-determining transcription factors prime cis-regulatory elements required for macrophage and B cell identities. *Mol Cell*. 2010;38(4):576-589.
15. Kharchenko PV, Tolstorukov MY, Park PJ. Design and analysis of ChIP-seq experiments for DNA-binding proteins. *Nat Biotechnol*. 2008;26(12):1351-1359.
16. Quinlan AR, Hall IM. BEDTools: a flexible suite of utilities for comparing genomic features. *Bioinformatics*. 2010;26(6):841-842.
17. Longabaugh WJR, Davidson EH, Bolouri H. Visualization, documentation, analysis, and communication of large-scale gene regulatory networks. *Biochimica et Biophysica Acta (BBA) - Gene Regulatory Mechanisms*. 2009;1789(4):363-374.
18. Zanini F, Berghuis BA, Jones RC, et al. Northstar enables automatic classification of known and novel cell types from tumor samples. *Sci Rep*. 2020;10(1):15251.
19. Setty M, Kiseliovas V, Levine J, Gayoso A, Mazutis L, Pe'er D. Characterization of cell fate probabilities in single-cell data with Palantir. *Nat Biotechnol*. 2019;37(4):451-460.

20. La Manno G, Soldatov R, Zeisel A, et al. RNA velocity of single cells. *Nature*. 2018;560(7719):494-498.
21. Bergen V, Lange M, Peidli S, Wolf FA, Theis FJ. Generalizing RNA velocity to transient cell states through dynamical modeling. *Nat Biotechnol*. 2020.
22. Kim D, Pertea G, Trapnell C, Pimentel H, Kelley R, Salzberg SL. TopHat2: accurate alignment of transcriptomes in the presence of insertions, deletions and gene fusions. *Genome Biol*. 2013;14(4):R36.
23. Love MI, Huber W, Anders S. Moderated estimation of fold change and dispersion for RNA-seq data with DESeq2. *Genome Biol*. 2014;15(12):550.
24. Subramanian A, Tamayo P, Mootha VK, et al. Gene set enrichment analysis: a knowledge-based approach for interpreting genome-wide expression profiles. *Proc Natl Acad Sci U S A*. 2005;102(43):15545-15550.
